# ABI5-FLZ13 Module Transcriptionally Represses Growth-related Genes to Delay Seed Germination in Response to ABA

**DOI:** 10.1101/2023.02.06.527385

**Authors:** Chao Yang, Xibao Li, Shunquan Chen, Chuanliang Liu, Lianming Yang, Kailin Li, Jun Liao, Xuanang Zheng, Hongbo Li, Yongqing Li, Shaohua Zeng, Xiaohong Zhuang, Pedro L. Rodriguez, Ming Luo, Ying Wang, Caiji Gao

## Abstract

The bZIP transcription factor ABSCISIC ACID INSENSITIVE5 (ABI5) is a master regulator determining seed germination and postgerminative growth in response to abscisic acid (ABA), but the detailed molecular mechanism underlying the repression function of ABI5 in plant growth remains to be characterized. In this study, we used proximity labeling (PL) to map the neighboring proteome of ABI5 and identified FCS-LIKE ZINC FINGER PROTEIN 13 (FLZ13) as an up-to-now unknown ABI5-interacting partner. Phenotypic analysis of *flz13* mutants and *FLZ13*-overexpressing lines demonstrated that FLZ13 acts as a positive regulator of ABA signaling. Interestingly, transcriptomic analysis showed that ABI5 and FLZ13 co-downregulate the expression of ABA-repressed and growth-related genes involved in chlorophyll biosynthesis, photosynthesis and cell wall organization, thereby repressing seed germination and seedling establishment in response to ABA. Further genetic analysis proved that FLZ13 works synergistically with ABI5 to regulate seed germination. Collectively, our study reveals a previously uncharacterized transcriptional regulatory mechanism regarding ABA-inhibited seed germination and seedling establishment.

## Introduction

Abscisic acid (ABA), a kind of sesquiterpenoid-based compound discovered in 1960s (Addicott et al., 1968; Cornforth et al., 1965; Eagles and Wareing, 1963; Ohkuma et al., 1963), is important for plant stress tolerance, growth and development (Chen et al., 2020; Cutler et al., 2010; Holdsworth et al., 2008; Leung and Giraudat, 1998). ABA is perceived by the soluble ABA receptors PYRABACTIN RESISTANCE1 (PYR1)/PYR1-LIKE (PYL)/REGULATORY COMPONENT OF ABA RECEPTOR (RCAR) (Ma et al., 2009; Nishimura et al., 2009; Park et al., 2009; Santiago et al., 2009). Upon ABA binding, these receptors can form a complex with TYPE 2C PROTEIN PHOSPHATASES (PP2Cs), thereby leading to the relief of inhibition of PP2Cs to the subfamily III of SUCROSE NONFERMENTING-1-RELATED PROTEIN KINASE 2s (SnRK2s) including SnRK2.2, SnRK2.3 and SnRK2.6 (Chen *et al*., 2020; Cutler *et al*., 2010; Ma *et al*., 2009; Miyazono et al., 2009; Nishimura *et al*., 2009; Park *et al*., 2009). Additionally, B2- and B3-type Raf protein kinases directly phosphorylate and activate subfamily III SnRK2s (Lin et al., 2021; Soma et al., 2020). Activated SnRK2s in turn phosphorylate and activate downstream transcription factors (TFs) such as ABA INSENSITIVE 5 (ABI5) and its homologs ABRE BINGDING FACTOR1/2/3/4 (ABF1/2/3/4), which subsequently modulate the expression of ABA-responsive genes through binding to the G-box motifs known as ABA response elements (ABREs) in their promoters (Fujii et al., 2007; Fujii and Zhu, 2009; Furihata et al., 2006; Kobayashi et al., 2005; Nakashima et al., 2009).

ABI5 as a bZIP transcription factor that is preferentially expressed in dry seeds and strongly induced by exogenous ABA, plays a critical role in ABA-mediated seed germination and postgerminative growth in *Arabidopsis* (Brocard et al., 2002; Finkelstein et al., 2005; Finkelstein, 1994; Finkelstein and Lynch, 2000; Lopez-Molina and Chua, 2000; Lopez-Molina et al., 2001; Lopez-Molina et al., 2002; Skubacz et al., 2016; Yu et al., 2015). The *ABI5* loss-of-function mutant plants are insensitivity to ABA, whereas plants overexpressing *ABI5* display hypersensitivity to ABA (Finkelstein, 1994; Finkelstein and Lynch, 2000; Lopez-Molina and Chua, 2000; Zhou et al., 2015). As a transcription factor, ABI5 functions mainly through modulating the expression of its target genes (Skubacz *et al*., 2016). Up to now, numerous ABI5-induced target genes have been identified (Lee and Luan, 2012; Skubacz *et al*., 2016), including *EARLY METHIONINE-LABELED 1* (*EM1*), *EM6* and *LEAD34* (Finkelstein and Lynch, 2000), which mainly regulate seed maturation but rarely perform functions in ABA-inhibited seed germination and postgerminative growth. Recently, Huang et al. reported that *PHOSPHATE1* (*PHO1*), an important regulator for phosphorus transport in *Arabidopsis* (Hamburger et al., 2002), is an ABA-repressed gene targeted by ABI5 to control seed germination (Huang et al., 2017). Although loss-of-function mutation in *PHO1* completely abolished the ABA-insensitive phenotype of *abi5-8* mutant (Zhou *et al*., 2015), the function of *PHO1* in ABA-inhibited seed germination remains elusive (Huang *et al*., 2017).

ABI5 often associates with other transcription regulators, such as ABI3, DELLA, BRI1-EMSSUPPRESSOR1 (BES1), JASMONATEZIM-DOMAIN (JAZ) proteins, VQ18/26, and INDUCER OF CBF EXPRESSION1 (ICE1) to regulate ABA responses (Hu et al., 2019; Ju et al., 2019; Lim et al., 2013; Pan et al., 2018; Zhao et al., 2019). Notably, most of the aforementioned interactors function as negative regulators to suppress the transcriptional regulation activity of ABI5, while the positive regulators of ABI5 are less well identified. In the case of ABI3, different results suggested the formation of molecular complexes with ABI5 to activate promoters containing ABREs, but other results also suggested that ABI5 can act as a transcription factor without ABI3 (Lopez-Molina *et al*., 2002). Thus, ABI5 can rescue the *abi3-1* mutant and acts downstream of ABI3 to induce growth arrest (Lopez-Molina *et al*., 2002). Recently, one study showed that the circadian clock proteins PSEUDO-RESPONSE REGULATOR 5 (PRR5) and PRR7 interact with and stimulate ABI5 to positively modulate ABA signaling during seed germination (Yang et al., 2021). Despite these findings, details of the transcriptional regulatory mechanisms underlying ABI5 and its cofactors still remain to be further characterized.

Previous studies reported that endospermic ABA represses the expression of cutin biosynthetic genes in the embryo through ABI5 action, which favors the ABA-mediated inhibition of the embryo-to-seedling transition (De Giorgi et al., 2015). In this study, we identified a hitherto unknown ABI5-interacting protein, FCS-LIKE ZINC FINGER PROTEIN 13 (FLZ13), which functions interdependently with ABI5 to determine the transcription of ABA-repressed and growth-related genes, thus positively modulates ABA response during seed germination and postgerminative growth.

## Results

### Proximity labeling proteomics identifies FLZ13 as a novel ABI5-interacting protein

Our recent study revealed that the plant-unique ESCRT component FYVE DOMAIN PROTEIN REQUIRED FOR ENDOSOMAL SORTING 1 (FREE1) physically interacts with ABI5 and interferes with the DNA binding ability of ABI5 to negatively regulate seedling establishment in response to ABA (Li et al., 2019; Li et al., 2022a). To further gain insights on the mechanism of ABI5-involed transcriptional regulation, we employed TurboID, a highly sensitive enzyme used in protein proximity labeling (Mair et al., 2019), to map the neighboring proteome of ABI5. To this end, we generated transgenic plants expressing either the ABI5-TurboID-GFP fusion or TurboID-GFP as a control in Col-0 wild type **(Figure 1A)**. The ABI5-TurboID-GFP fusion is functional in planta, because ABI5-TurboID-GFP correctly localized to the nucleus and the transgenic plants were hypersensitive to ABA treatment **(Supplemental Figure 1)**. 5-d-old seedlings were treated with 50 μM biotin for 1 h, then harvested to isolate total proteins for affinity purification with streptavidin-coated beads followed by label-free quantitative mass spectrometry (LFQMS). Using *P* value < 0.05 and fold change > 2 as cutoffs, we obtained 67 ABI5-specific preys **(Figure 1A; Supplemental Data 1)**. Among them, ABI5 BINDING PROTEIN1 (AFP1) and FREE1 are previously reported as ABI5-interacting proteins (Li *et al*., 2019; Lopez-Molina *et al*., 2003), indicating that our TurboID-based proximity labeling is effective to isolate ABI5 interactors. To gain an overview of these candidates, 48 nuclear-localized proteins out of the 67 higher enriched proteins were used to construct a protein-protein interaction (PPI) network **(Supplemental Data 2)**. As shown in **Supplemental Figure 2A**, 44 out of 48 nuclear proteins were included in this network, indicating that they were indeed associated in plant cells. To enrich the ABI5-interacting network, we generated a bigger network combinedly using the 48 nuclear proteins identified in this study and 37 known ABI5-interacting proteins **(Supplemental Data 3)**. Finally, 82 out of 83 proteins could be connected in this network **(Supplemental Figure 2B)**, indicating that ABI5 can associate with diverse proteins to generate multiple functional complexes and that TurboID-mediated PL-LFQMS is a powerful approach to identify protein interactome in plants.

**Figure 1:**
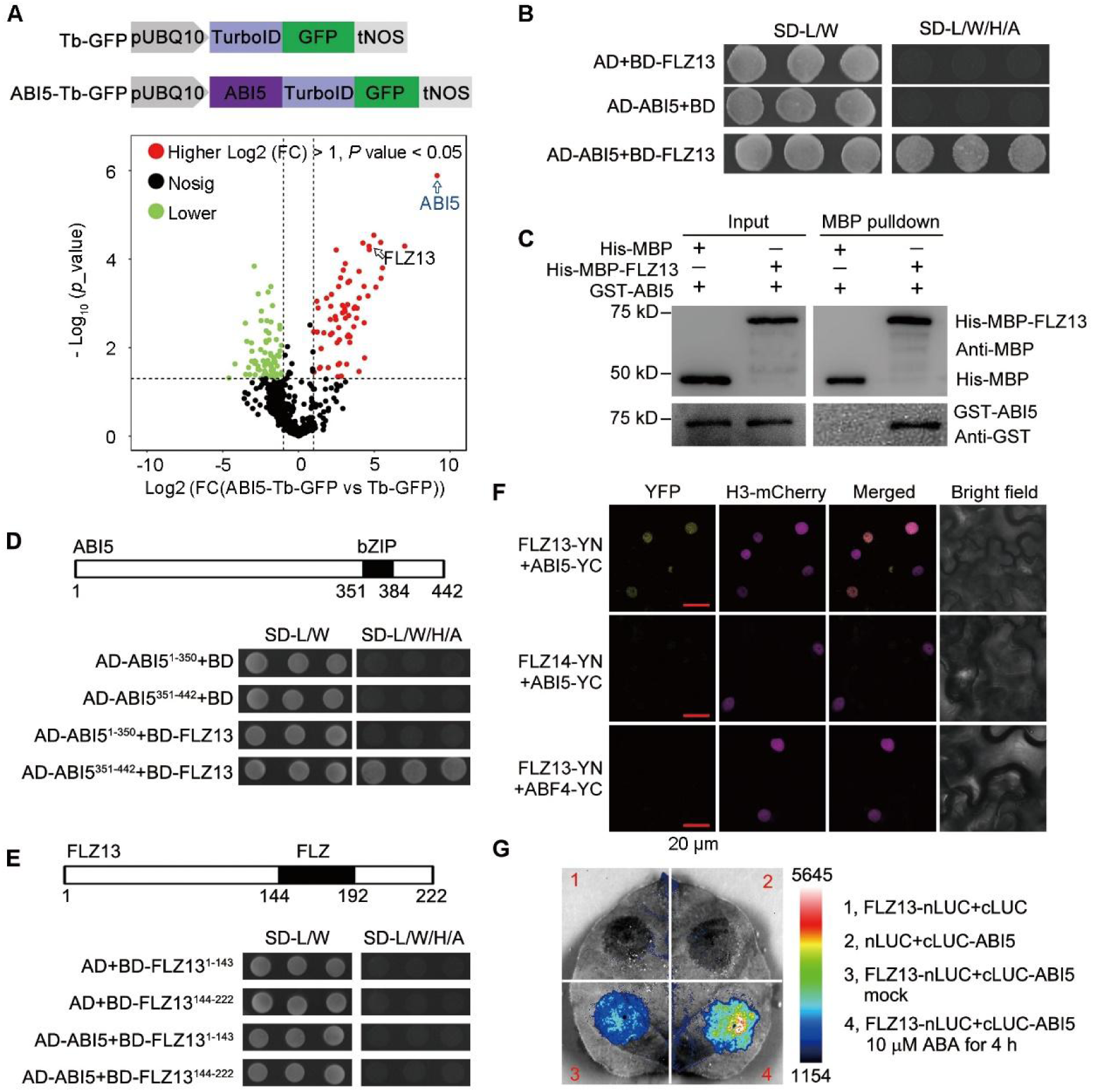
Identification of FLZ13 as a novel interactor of ABI5. A, Volcano plot showing protein abundance in ABI5-Tb-GFP vs Tb-GFP pair. The constructs encoding ABI5-Tb-GFP and Tb-GFP fusion proteins were shown. The integrated LFQ (label-free quantitation) peptide intensity data from three biological replicates (*n* = 3 biologically independent proximity labeling experiments) of each sample were normalized and subjected to ratiometric analysis and plotting by using Perseus program. In total, 611 proteins identified in both ABI5-Tb-GFP and Tb-GFP samples were plotted. Preys with Preys with fold change > 2 (Log2 fold change > 1) and *p* < 0.05 (-Log10 *P* value > 1.301) were marked by colored dots. Green and red dots represent significantly decreased and increased proteins, respectively. B, Y2H assay showing the interaction between ABI5 and FLZ13. Protein interaction was determined by growth of the yeast cells co-transformed with various combinations of the plasmids on synthetic dropout medium lacking Leu and Trp (SD-L/W) and synthetic dropout medium lacking Leu, Trp, His, and adenine (SD-L/W/H/A). C, *In vitro* assay showing the physical binding of ABI5 and FLZ13. Equal amount GST-ABI5 were incubated with His-MBP and His-MBP-FLZ13 bound to MBP magnetic beads and then eluted and analyzed by immunoblotting using Anti-GST and anti-MBP antibodies. D-E, Mapping the interaction region of ABI5 and FLZ13 by Y2H. Diagrams of ABI5 and FLZ13 protein structure were shown. Protein interaction was determined by growth of the yeast cells co-transformed with various combinations of the plasmids on synthetic dropout medium lacking Leu and Trp (SD-L/W) and synthetic dropout medium lacking Leu, Trp, His, and adenine (SD-L/W/H/A). F, BiFC assay revealing the association of ABI5 and FLZ13 in the nucleus in *N. benthamiana* leaves. FLZ13 fused to the N-terminal half of YFP (FLZ13-YN) was co-infiltrated with ABI5 fused to the C-terminal half of YFP (ABI5-YC). The FLZ13-YN/ABF4-YC and ABI5-YC/FLZ14-YN pairs were used as negative controls. G, LCI assay showing the interaction between ABI5 and FLZ13 in *N. benthamiana* leaves. Agrobacteria harboring FLZ13-nLUC was co-infiltrated with agrobacteria harboring cLUC-ABI5 into *N. benthamiana* leaves. The FLZ13-nLUC/cLUC and cLUC-ABI5/nLUC pairs were used as negative controls. After 36 h co-infiltration, the leaves were treated with mock and 10 μM ABA for 4 h followed by luminescence imaging.

Among the identified proteins in this study, FCS-LIKE ZINC FINGER PROTEIN 13 (FLZ13) is the most interesting one, since its gene expression is induced by ABA **(Supplemental Data 2)** and its molecular function has not been characterized yet. Therefore, we chose FLZ13 as an example for characterizing its function in ABI5-mediated signaling pathway. Yeast-two-hybrid (Y2H) analysis showed that FLZ13 could specifically interact with ABI5 but not with other transcription factors in ABA signaling including ABI3, ABI4, ABF2 and ABF4 **(Figure 2B; Supplemental Figure 3)**. The direct interaction between FLZ13 and ABI5 was further corroborated by *in vitro* pull-down assay, which demonstrated that GST-ABI5 could be specifically precipitated by His-MBP-FLZ13 but not by His-MBP alone **(Figure 1C)**. Additional Y2H assays with truncated proteins showed that the C-terminal part of ABI5 (351-442 aa) harboring a bZIP domain but not the N-terminal part of ABI5 (1-350 aa) was able to bind FLZ13 **(Figure 1D)**. Furthermore, we found that truncated FLZ13 lacking of either the N-terminal or the C-terminal half failed to interact with ABI5 **(Figure 1E)**. These Y2H results suggested that the C-terminal bZIP domain of ABI5 mediates interaction with the full-length FLZ13. To confirm the interaction of ABI5 and FLZ13 in planta, we performed bimolecular fluorescence complementation (BiFC) and luciferase complementation imaging (LCI) assays. The obtained BiFC results revealed that ABI5 interacted with FLZ13 in the nucleus **(Figure 1F)**. Furthermore, co-expression of FLZ13-nLUC with cLUC-ABI5 in tobacco leaves resulted in obvious LUC signals **(Figure 1G; Supplemental Figure 4)**, whose activity could be significantly boosted by ABA treatment, indicating that ABA might promote the formation of ABI5-FLZ13 complex **(Figure 1G; Supplemental Figure 4)**. Collectively, these results clearly demonstrated that FLZ13 physically interacts with ABI5, suggesting that FLZ13 might function as a working partner of ABI5 to mediate ABA signaling.

**Figure 2:**
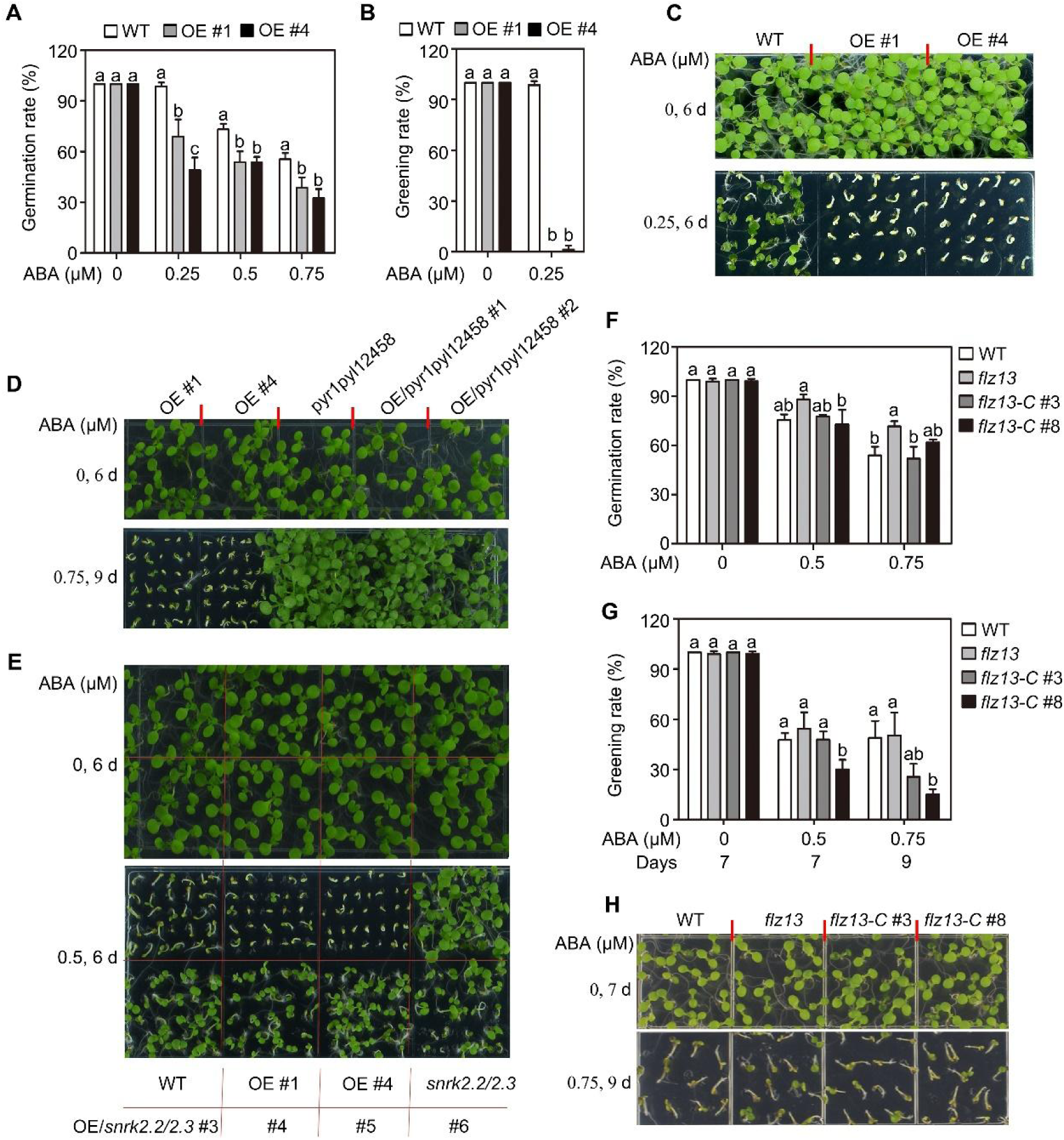
FLZ13 positively regulates ABA signaling during seed germination. A, Germination of WT and two *FLZ13*-overexpression lines (OE #1 and OE #4). Seed germination was recorded 4 d after germination on 1/2 MS medium supplemented with different concentrations of ABA. Data are presented as means ± SD (*n* = 3 biological replicates). For each biological replicate, around 60 seeds of each genotype were used for the germination assay. The different letters above each bar indicate statistically significant differences as determined by a one-way ANOVA test followed by Tukey’s multiple test (P < 0.05). B, Cotyledon greening of WT and *FLZ13*-OE lines. Cotyledon greening was scored 6 d after germination on 1/2 MS medium supplemented with 0 or 0.25 μM ABA. Data are presented as means ± SD (*n* = 3 biological replicates). For each biological replicate, around 60 seeds of each genotype were used for seedling establishment assay. The different letters above each bar indicate statistically significant differences as determined by a one-way ANOVA test followed by Tukey’s multiple test (P < 0.05). C, Photographs of seedlings of WT and two *FLZ13*-OE lines germinated on medium containing 0 or 0.25 μM ABA for 6 d. D, Seedlings of *FLZ13-OE*, *pyr1pyl12458*, and *FLZ13*-OE/*pyr1pyl12458* upon 0 μM ABA for 6 d or 0.75 μM ABA treatment for 9 d. E, Seedlings of WT, *FLZ13*-OE, *snrk2.2*/*2.3*, and *FLZ13*-OE/*snrk2.2*/*2.3* upon 0 or 0.5 μM ABA treatment for 6 d. F, Quantification of germination rates of WT, *flz13* and two complementary lines. Seed germination was recorded 4 d after germination on 1/2 MS medium supplemented with different concentrations of ABA. Data are presented as means ± SD (*n* = 3 biological replicates). For each biological replicate, around 60 seeds of each genotype were used for the germination assay. The different letters above each bar indicate statistically significant differences as determined by a one-way ANOVA test followed by Tukey’s multiple test (P < 0.05). G, Quantification of cotyledon greening rates of WT, *flz13* and two complementary lines treated with different concentrations of ABA as recorded at the indicated time points. Data are presented as means ± SD (*n* = 3 biological replicates). For each biological replicate, around 60 seeds of each genotype were used for seedling establishment assay. The different letters above each bar indicate statistically significant differences as determined by a one-way ANOVA test followed by Tukey’s multiple test (P < 0.05). H, Photographs of seedlings of WT, *flz13* and two complementary lines upon 0 μM ABA for 7 d or 0.75 μM ABA treatment for 9 d.

### *FLZ13* is responsive to ABA in germinating seeds and positively regulates ABA response

Given that FLZ13 is a novel interactor of ABI5, we first examined the gene and protein expression of *FLZ13* during seed germination with or without exogenous ABA treatment. Consistent with previous reports (Brocard *et al*., 2002; Lopez-Molina *et al*., 2001), the expression of *ABI5* decreased substantially during seed germination, but increased upon ABA treatment **(Supplemental Figure 5A** and **B)**. However, *FLZ13* displayed slightly increased expression pattern during seed germination or upon ABA treatment **(Supplemental Figure 5A** and **B)**. To determine the protein abundance of FLZ13 in germinating seeds, transgenic *Arabidopsis* lines expressing a *pFLZ13::FLZ13-GFP* fusion in Col-0 background were generated. As presented in **Supplemental Figure 5C**, ABI5 protein showed a remarkably decreased expression, while the FLZ13-GFP protein displayed a gradual accumulation following seed germination. When germinating seeds were exposed to ABA, the protein level of FLZ13-GFP had not obvious change, but that of ABI5 was increased acutely **(Supplemental Figure 5C)**. These results indicated the co-existence of FLZ13 with ABI5 in the presence of ABA, meeting the requirement for their interaction in planta.

The biological function of *FLZ13* in *Arabidopsis* has not been documented yet. Since FLZ13 physically interacts with ABI5 **(Figure 1)** and responds to ABA in germinating seed **(Supplemental Figure 5)**, we hypothesized that FLZ13 might be involved in ABI5-mediated ABA signaling during seed germination. To test this possibility, we first generated *FLZ13* overexpression (OE) transgenic plants in Col-0 background and two representative T3 lines (*pUBQ10::FLZ13-GFP #1* and *pUBQ10::FLZ13-GFP #4,* termed as *FLZ13-*OE #1 and OE #4) with higher gene expression levels under normal growth conditions were chosen for further study **(Supplemental Figure 6)**. As shown in **Figure 2A-C**, the seed germination and seedling growth of WT and *FLZ13*-OE lines did not show obvious differences in the medium without exogenous ABA, whereas the seeds of *FLZ13*-OE lines had much lower germination and greening rates than Col-0 in the presence of ABA, suggesting that overexpression of *FLZ13* conferred ABA hypersensitivities of seed germination and seedling establishment. We also overexpressed *FLZ13* in the *pyr1pyl12458* and *snrk2.2/2.3* background for genetic analysis of FLZ13 and other components involved in ABA signaling (Fujii *et al*., 2007; Gonzalez-Guzman et al., 2012). The obtained results of ABA sensitivity assay showed that *FLZ13*-OE/*pyr1pyl12458* and *FLZ13*-OE/*snrk2.2*/*2.3* lines phenocopied the *pyr1pyl12458* and *snrk2.2*/*2.3* mutant, respectively **(Figure 2D** and **E)**, suggesting that the effect of *FLZ13* overexpression in conferring ABA hypersensitivity requires a functional ABA signaling pathway.

To further confirm the roles of *FLZ13* in ABA signaling, we obtained a *FLZ13* T-DNA insertion mutant (SALK_142112C) harboring a T-DNA insertion at 100 bp upstream of 5’ untranslated region (UTR), which results in an obvious reduction (20% of wild-type expression) in the transcription level of *FLZ13* **(Supplemental Figure 6)**. The ABA-sensitivity test showed that *flz13* mutant was slightly more resistant to ABA than Col-0 during, in terms of seed germination rate **(Figure 2F** and **3A)**. We next introduced a construct containing the entire genomic sequence of *FLZ13* into the *flz13* mutant **(Supplemental Figure 6)** resulting in 1.9-fold (*gFLZ13-C3*) and 2.38-flod (*gFLZ13-C8*) expression of *FLZ13*, respectively. Phenotype analysis showed that these two complementation lines germinated like the WT plants in response to ABA **(Figure 2F)**, but with lower greening rates than WT and *flz13* **(Figure 2G** and **H)**, suggesting that the reduced ABA sensitivity of *flz13* mutant is indeed because of the knockdown of *FLZ13*. To obtain additional mutant alleles, we used CRISPR/Cas9 technology to edit *FLZ13* in Col-0 wild-type plants. We obtained one edited line, termed *flz13-cas9 #2*, with a deletion of 16 nucleotides (164 to 179) that results in a predicted truncated FLZ13 protein **(Supplemental Figure 6A)**. Similar to *flz13* **(Figure 3B)**, the *flz13-cas9 #2* mutant showed a reduced sensitivity to ABA as revealed by comparison the greening rate in the presence of 1.0 µM ABA **(Supplemental Figure 6B)**. Taken together, these data suggest that FLZ13 positively regulates ABA signaling to repress seed germination and early seedling growth in *Arabidopsis*.

**Figure 3:**
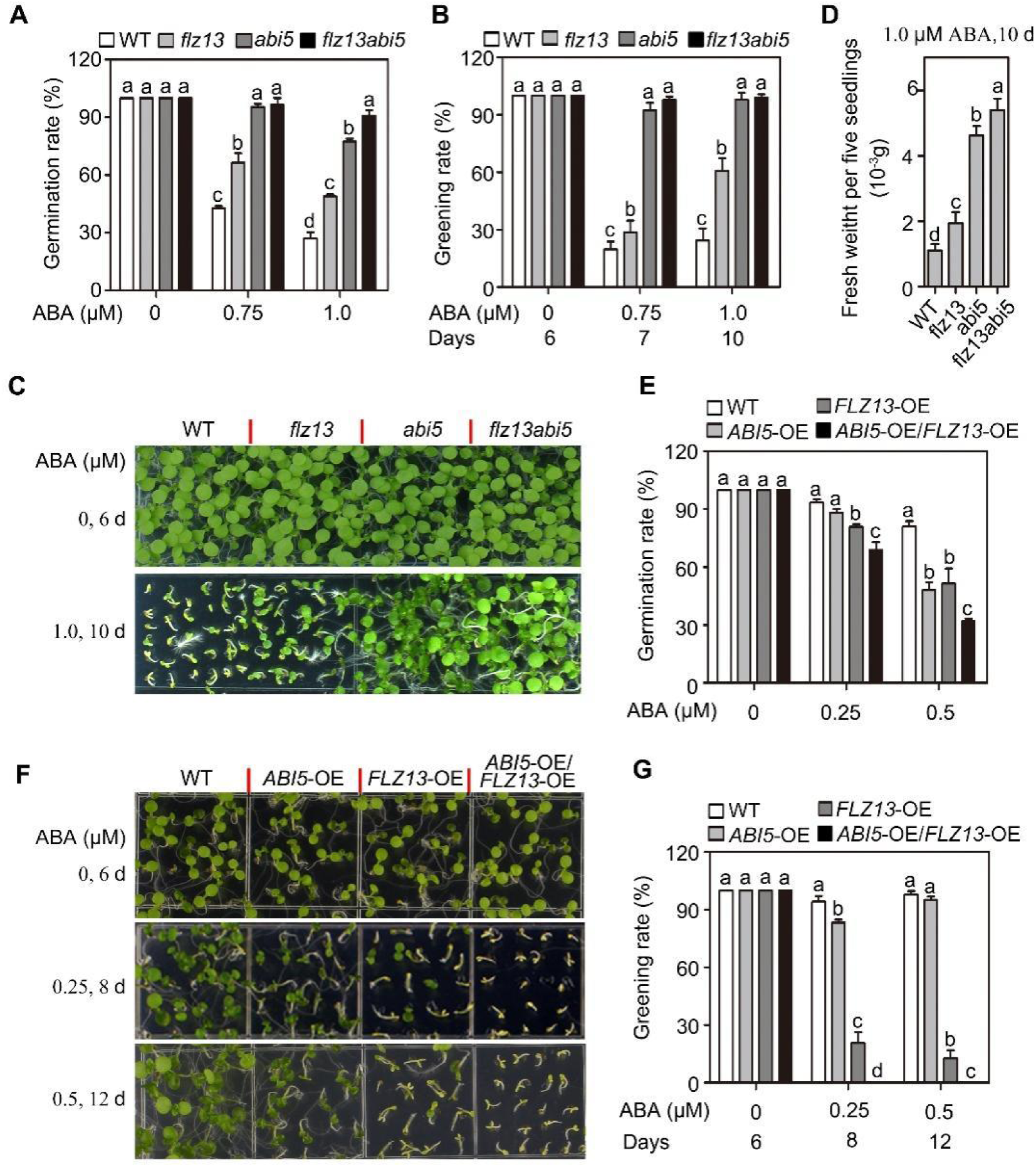
Genetic interaction between FLZ13 and ABI5 in ABA signaling during seed germination. A, Germination of WT, *flz13*, *abi5-8* and *flz13 abi5-8* mutants treated with different concentrations of ABA. The germination rate was recorded 4 d after planting on 1/2 MS medium. Data are presented as means ± SD (*n* = 3 biological replicates). For each biological replicate, around 60 seeds of each genotype were used for the germination assay. The different letters above each bar indicate statistically significant differences as determined by a one-way ANOVA test followed by Tukey’s multiple test (P < 0.05). B, Cotyledon greening of WT, *flz13*, *abi5-8* and *flz13 abi5-8* treated with different concentrations of ABA as recorded at the indicated time points. Data are presented as means ± SD (*n* = 3 biological replicates). For each biological replicate, around 60 seeds of each genotype were used for seedling establishment assay. The different letters above each bar indicate statistically significant differences as determined by a one-way ANOVA test followed by Tukey’s multiple test (P < 0.05). C, Seedlings of WT, *flz13*, *abi5-8* and *flz13 abi5-8* treated with different concentrations of ABA for the indicated time points. D, Fresh weight of five seedlings of WT, *flz13*, *abi5-8* and *flz13 abi5-8* treated with 1.0 μM ABA for 10 d. Data are presented as means ± SD (*n* = 5 biological replicates). The different letters above each bar indicate statistically significant differences as determined by a one-way ANOVA test followed by Tukey’s multiple test (P < 0.05). E, Germination of WT, *ABI5*-OE, *FLZ13-OE* and *ABI5*-OE/*FLZ13*-OE treated with different concentrations of ABA. The germination rate was recorded 4 d after planting on 1/2 MS medium. Data are presented as means ± SD (*n* = 3 biological replicates). For each biological replicate, around 60 seeds of each genotype were used for the germination assay. The different letters above each bar indicate statistically significant differences as determined by a one-way ANOVA test followed by Tukey’s multiple test (P < 0.05). F, Seedlings of WT, *ABI5*-OE, *FLZ13-OE* and *ABI5*-OE/*FLZ13*-OE germinated on medium containing different concentrations of ABA for the indicated time points. G, Cotyledon greening of WT, *ABI5*-OE, *FLZ13-OE* and *ABI5*-OE/*FLZ13*-OE treated with different concentrations of ABA as recorded at the indicated time points. Data are presented as means ± SD (*n* = 3 biological replicates). For each biological replicate, around 60 seeds of each genotype were used for seedling establishment assay. The different letters above each bar indicate statistically significant differences as determined by a one-way ANOVA test followed by Tukey’s multiple test (P < 0.05).

### FLZ13 functions synergistically with ABI5 to modulate ABA signaling

To determine the genetic relationship between ABI5 and FLZ13 in ABA signaling, we crossed *flz13* with *abi5-8* to generate a *flz13 abi5-8* double mutant for phenotypic analysis. Consistent with previous studies (Zhou *et al*., 2015), the *flz13* and *abi5-8* seeds germinated and grew faster than Col-0 seeds on ABA-containing medium **(Figure 3A-D)**. Notably, the *flz13 abi5-8* double mutant displayed less sensitivity to ABA treatment in comparison with the *abi5-8* and *flz13* single mutants when growing on the medium containing 1.0 µM ABA **(Figure 3A-D)**, suggesting that FLZ13 and ABI5 may function synergistically to regulate ABA signaling during seed germination. We also crossed our previously established *pUBQ10::ABI5-GFP #1* (*ABI*5-OE *#1*, Li et al., 2019) with *FLZ13*-OE *#1* to generate plants simultaneously overexpressing *ABI5* and *FLZ1*3 (*ABI5*-OE/*FLZ13*-OE) for testing its ABA sensitivity. As expected, the progenies of *ABI5*-OE/*FLZ13*-OE displayed significantly lower rates of seed germination and cotyledon greening than those of *ABI5*-OE and *FLZ13*-OE in the medium with various concentrations of ABA **(Figure 3E-G)**, suggesting an additive effect of ABI5 and FLZ13 in plant response to ABA. Collectively, these data proved that FLZ13 and ABI5 functions synergistically to repress seed germination upon ABA treatment.

### FLZ13 and ABI5 co-regulate common target genes in response to ABA

The above observation of the protein and genetic interactions between FLZ13 and ABI5 motivated us to identify FLZ13/ABI5 co-regulated genes in response to ABA during seed germination. RNA-seq analysis was carried out using germinating seeds of WT, *ABI5*-OE *#1* and *FLZ1*3-OE #1 treated with 0.5 μM ABA. In WT seeds, 6970 deferentially expressed genes (DEGs) (fold change ≥ 2, False Discovery Rate (FDR) < 0.01) were identified **(Figure 4A; Supplemental Data 4)**, including 77 genes previously found to be induced by ABA **(Supplemental Data 5)**, indicating that this batch of RNA-seq data was reliable. Compared to WT, 1161 and 1181 DEGs were respectively identified in *ABI5*-OE *#1* and *FLZ1*3-OE #1 seeds with 0.5 μM ABA treatement **(Figure 4A; Supplemental Data 6 and 7)**.

**Figure 4:**
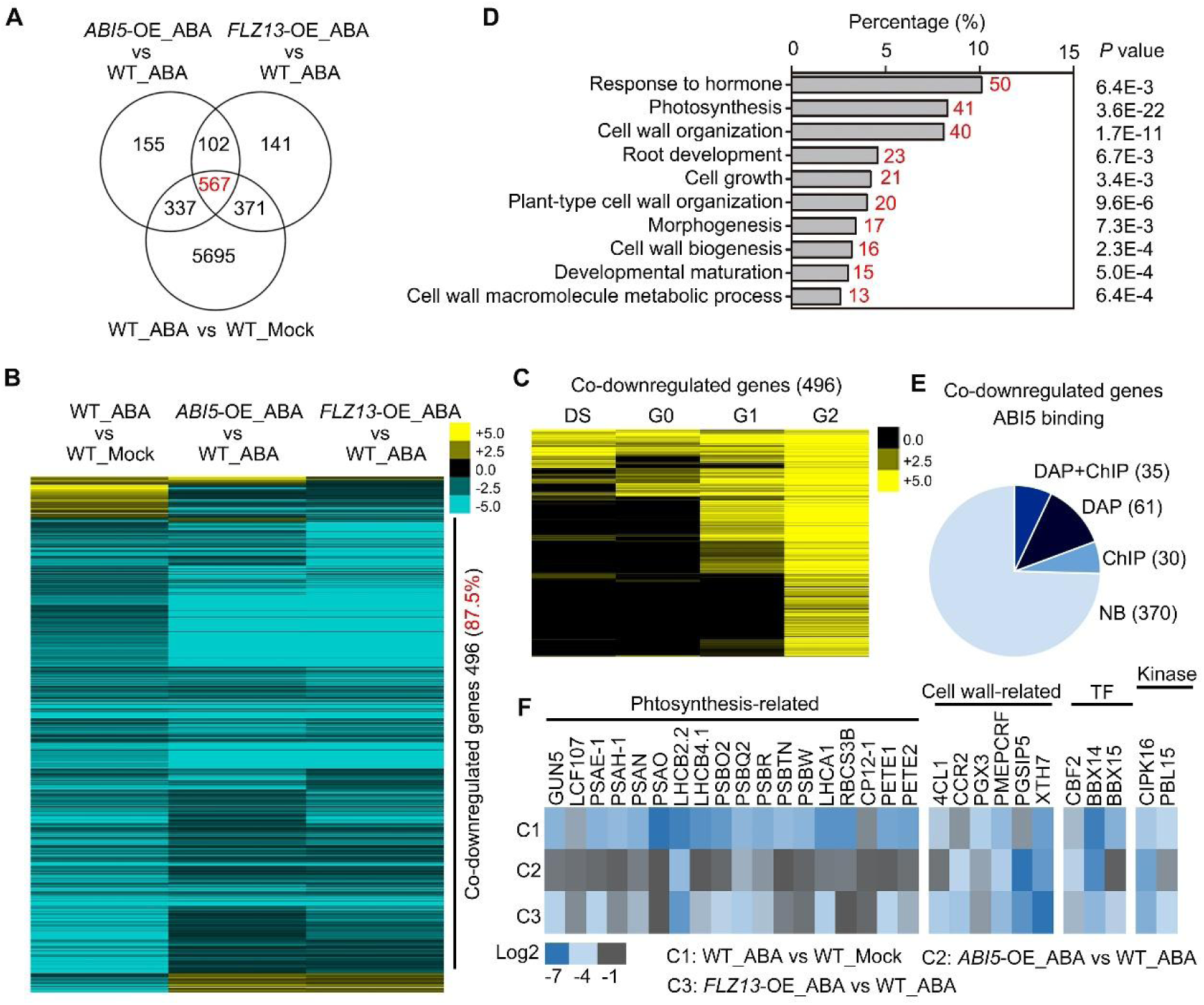
Identification of ABA/ABI5/FLZ13-co-regulated genes. A, Venn diagram showing the overlap of ABA/ABI5/FLZ13-regulated genes. B, Heat map showing the expression of ABA/ABI5/FLZ13 co-regulated genes. The expression data were clustered by Cluster 3.0 software and edited in Tree-view. C, Heat map showing the expression of ABA/ABI5/FLZ13 co-repressed genes during seed germination. The expression data were obtained from publicly available database (http://bioinfo.sibs.ac.cn/plant-regulomics/; Ran et al., 2020) and clustered by Cluster 3.0 software and edited in Tree-view. DS, dry seed; G0, G1 and G2 means germinating seeds of 0 d, 1 d and 2 d, respectively. D, Representative enriched BP terms of ABA/ABI5/FLZ13 co-repressed genes. The GO enrichment analysis was performed by using online program (https://david.ncifcrf.gov/home.jsp; Huang et al., 2009). E, Pie chart showing the number of ABA/ABI5/FLZ13 co-repressed genes which are bound by ABI5 in ChIP-seq and/ or DAP-seq assay (C. O’Malley et al., 2016). NB, no bound. F, Heat map showing the expression of the well-annotated ABI5 target genes which are co-repressed by ABA/ABI5/FLZ13.

To identify ABI5 and FLZ13 co-regulated and ABA-responsive genes, we selected those genes that would meet the following three criteria: (i) genes that were differentially expressed after ABA treatment in Col-0; (ii) their expression differed between Col-0 and *ABI5*-OE #1 after ABA treatment; (iii) their expression differed between Col-0 and *FLZ1*3-OE #1 after ABA treatment. Finally, 567 genes were identified as ABA-responsive and ABI5-FLZ13 co-regulated genes **(Figure 4A, Supplemental Data 8)**. The heat map showed that only 4 genes were co-upregulated, while most (496, 87.5%) genes were simultaneously suppressed by ABA, ABI5 and FLZ13 **(Figure 4B)**, and they all showed induced expression pattern during seed germination **(Figure 4C)**. GO term enrichment analysis revealed that these co-repressed DEGs were mainly enriched in categories related to growth, including “photosynthesis” (41 genes, *p* value = 3.6E-22) and “cell wall organization” (40 genes, *p* value = 1.7E-11) **(Figure 4D, Supplemental Data 9)**. Furthermore, among the 496 genes, 126 genes were targeted by ABI5 as verified by ChIP-seq and/or DAP-seq assays (O’Malley *et al*., 2016) **(Figure 4E; Supplemental Data 10)**. The set of 35 highly confident ABI5-targeted genes, as verified by both ChIP-seq and DAP-seq assays, contains 15 photosynthesis-related genes, 6 cell wall modification-related genes, 3 transcription factors and 2 kinases **(Figure 4E, Supplemental Data 11)**. Collectively, these results confirmed that FLZ13 and ABI5 functions synergistically to regulate plant response to ABA by co-targeting a large number of ABA-responsive genes, especially ABA-repressed and growth-related genes.

### FLZ13 activation of ABA signaling requires a functional ABI5

To study the genetic epistasis between *FLZ13* and *ABI5*, we explored whether the action of FLZ13 in mediating ABA signaling requires a functional ABI5 by generating plants with overexpression of *FLZ1*3 in *abi5-8* mutant background. Progenies of *FLZ1*3-OE *#1*/*abi5-8* were hyposensitive to 0.5 μM ABA during seed germination with higher percentages of seed germination and cotyledon greening rates when compared with those of Col-0 and *FLZ1*3-OE *#1* plants **(Figure 5A-C)**. However, when compared with the *abi5-8,* the *FLZ1*3-OE *#1/abi5-8* was slightly more sensitive to ABA at 0.75 μM ABA, indicating that ABA hypersensitivity conferred by *FLZ1*3 overexpression largely but not fully depends on ABI5 **(Figure 5A-C)**.

**Figure 5:**
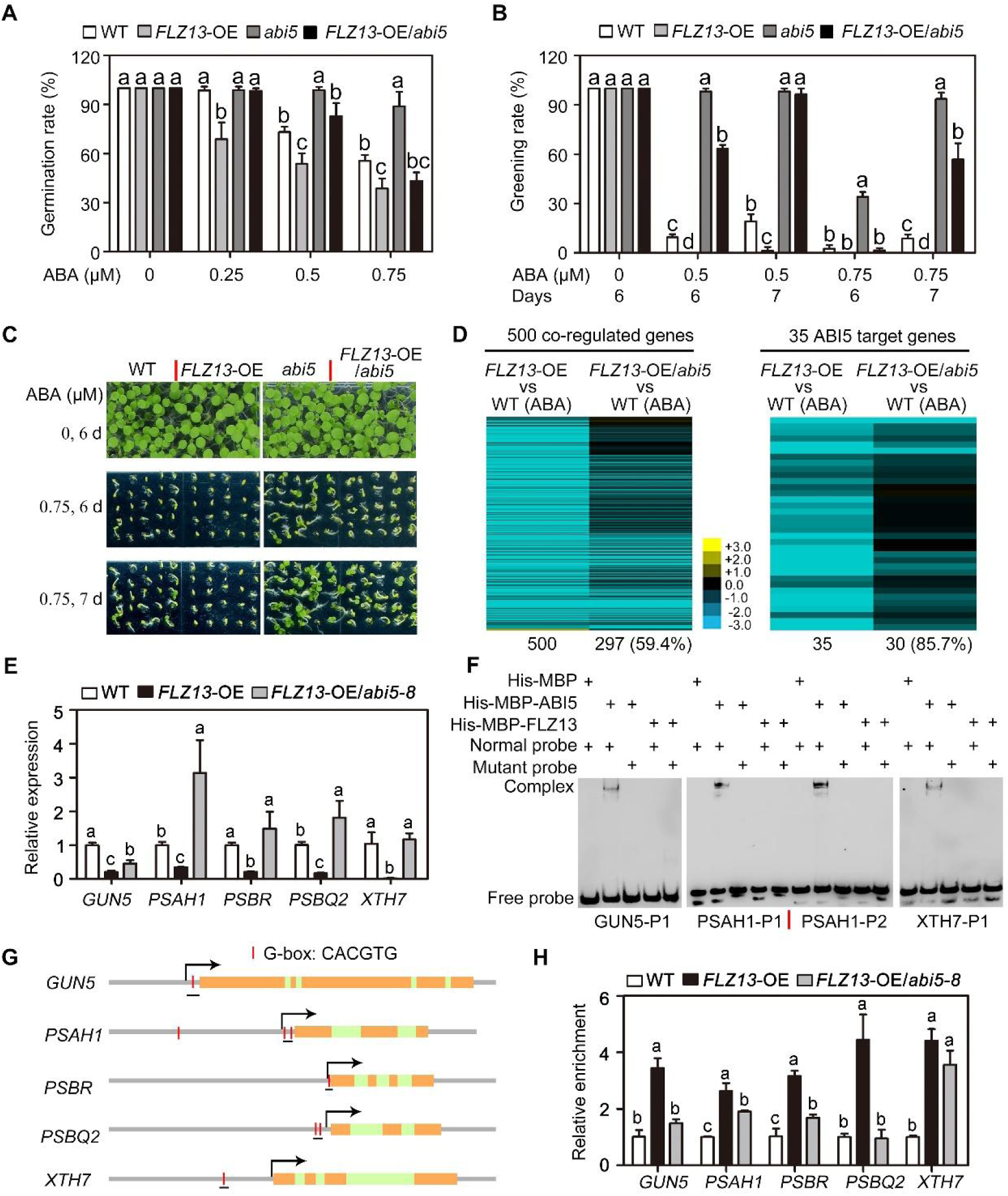
FLZ13 regulates ABA response during seed germination in an ABI5 dependent manner. A, Germination rate of WT, *FLZ13*-OE, *abi5-8* and *FLZ13*-OE/*abi5-8* treated with different concentrations of ABA. The germination rate was recorded 4 d after planting on 1/2 MS medium. Data are presented as means ± SD (*n* = 3 biological replicates). For each biological replicate, around 60 seeds of each genotype were used for the germination assay. The different letters above each bar indicate statistically significant differences as determined by a one-way ANOVA test followed by Tukey’s multiple test (P < 0.05). B, Cotyledon greening of WT, *FLZ13*-OE, *abi5-8* and *FLZ13*-OE/*abi5-8* treated with different concentrations of ABA for the indicated time points. Data are presented as means ± SD (*n* = 3 biological replicates). For each biological replicate, around 60 seeds of each genotype were used for seedling establishment assay. The different letters above each bar indicate statistically significant differences as determined by a one-way ANOVA test followed by Tukey’s multiple test (P < 0.05). C, Seedlings of WT, *FLZ13*-OE, *abi5-8* and *FLZ13*-OE/*abi5-8* germinated on the medium containing different concentrations of ABA for the indicated time points. D, Heat map showing the expression of ABA/ABI5/FLZ13 co-regulated genes in *FLZ13*-OE and *FLZ13*-OE/*abi5-8* lines in the present of 0.5 μM ABA. 3-day-old germinating seeds with 0.5 μM ABA treatments were collected for RNA-seq assay. E, qRT-PCR assay showing the expression of the selected genes in WT, *FLZ13*-OE and *FLZ13*-OE/*abi5-8* upon 0.5 μM ABA treatment. 3-day-old germinating seeds with 0.5 μM ABA treatments were collected for qRT-PCR assay. *PP2A* was used as the internal control. Data are presented as means ± SD (*n* = 3 technical replicates). The different letters above each bar indicate statistically significant differences as determined by a one-way ANOVA test followed by Tukey’s multiple test (P < 0.05). This experiment repeated twice with similar results. F, EMSA showing the binding of ABI5 and FLZ13 to the three ABI5 target genes *GUN5*, *PSAH1* and *XTH7*. The DNA probe were marked in Supplemental Figure 8. G, Gene structure of the selected ABI5 target genes. The position of G-box and the fragments used for ChIP-PCR assay were marked. H, ChIP-qPCR showing the relative enrichment of FLZ13 to the selected gene locus. Data are presented as means ± SD (*n* = 3 technical replicates). The different letters above each bar indicate statistically significant differences as determined by a one-way ANOVA test followed by Tukey’s multiple test (P < 0.05). This experiment repeated twice with similar results.

To assess the effect of ABI5 on the expression of *FLZ13-*regulated genes, we carried out RNA-seq analysis using germinating *FLZ1*3-OE *#1*/*abi5-8* seeds with ABA treatment **(Supplemental Data 12)**. Heat map of the 500 ABA/ABI5/FLZ13 synchronously co-regulated genes showed changed expression level and 297 out of these 500 genes (59.4%) exhibited significant alteration in *FLZ1*3-OE *#1/abi5-8* **(Figure 5D, leaf panel)**, supporting the notion that FLZ13-mediated transcription regulation largely depends on the function of ABI5. For example, 30 out of 35 ABI5 target genes showed differential expression between *FLZ1*3-OE *#1* and *FLZ1*3-OE *#1*/*abi5-8* plants after ABA treatment **(Figure 5D, right panel)**. We also conducted qRT-PCR to confirm the transcript levels of five ABI5 target genes (*GUN5*, *PSAH1*, *PSBR*, *PSBQ2* and *XTH7)* (**Supplemental Figure 7**). The obtained data revealed that mutation of ABI5 largely compromised *FLZ13*-repressed expression of these genes **(Figure 5E)**. The above ABA-sensitivity test and gene expression analysis clearly demonstrated that the function of FLZ13 in ABA signaling largely depends on ABI5.

To further explore how ABI5 affects the action of FLZ13, we employed electrophoretic mobility shift assay (EMSA) to examine the DNA binding ability of ABI5 and FLZ13 to their co-regulated genes. As shown in **Figure 5F**, ABI5 directly bound to the promoter regions of *GUN5*, *PSAH1*, and *XTH7* via a conserved G-box *ci*s-element, whereas FLZ13 failed to bind to them **(Supplemental Figure 7)**. To check these bindings *in vivo*, chromatin immunoprecipitation (ChIP) was performed using ABA-treated germinating seeds of *FLZ1*3-OE *#1* and *FLZ1*3-OE *#1*/*abi5-8* plants. ChIP-quantitative PCR (ChIP-qPCR) analysis showed that FLZ13 could be obviously enriched at the ABI5-bound genomic loci including *GUN5*, *PSAH1*, *PBSR*, *PBSQ2* and *XTH7* in the *FLZ1*3-OE *#1* plant **(Figure 5G and H)**. However, the enrichment of FLZ13 on their promoters was generally decreased in *FLZ1*3-OE *#1/abi5-8* than *FLZ1*3-OE *#1* **(Figure 5G and H)**. These findings imply that the recruitment of FLZ13 to the promoters of *GUN5*, *PSAH1*, *PBSR*, *PBSQ2* and *XTH7* at least partially depends on ABI5.

### Knockdown of *FLZ13* compromises the function of ABI5

To elucidate the roles of FLZ13 on the activity of ABI5, we generated a *ABI5*-OE/*flz13* plant by crossing and performed the ABA-sensitivity test. As shown in **Figure 6A and B**, knockdown of *FLZ13* also significantly elevated the greening rate of cotyledons of *ABI5*-OE #1 plants in the presence of ABA, suggesting that FLZ13 contributes to the ABI5-modulated plant ABA response. To explore how FLZ13 functions in ABI5-mediated ABA response, we first analyzed whether FLZ13 affects ABI5 protein stability and phosphorylation, two decisive factors contributing to ABI5 activity during ABA signaling (Yu et al., 2015). The obtained immunoblotting results demonstrated that the abundance of ABI5 protein was similar in *flz13* and Col-0 both under normal growth conditions and ABA treatment **(Figure 6C)**, suggesting that FLZ13 did not regulate the stability of ABI5. Moreover, the phosphorylation level of ABI5 was similar in both *flz13* and Col-0 plants with ABA treatment **(Figure 6D)**, suggesting that FLZ13 did not affect the phosphorylation of ABI5 in the presence of ABA.

**Figure 6:**
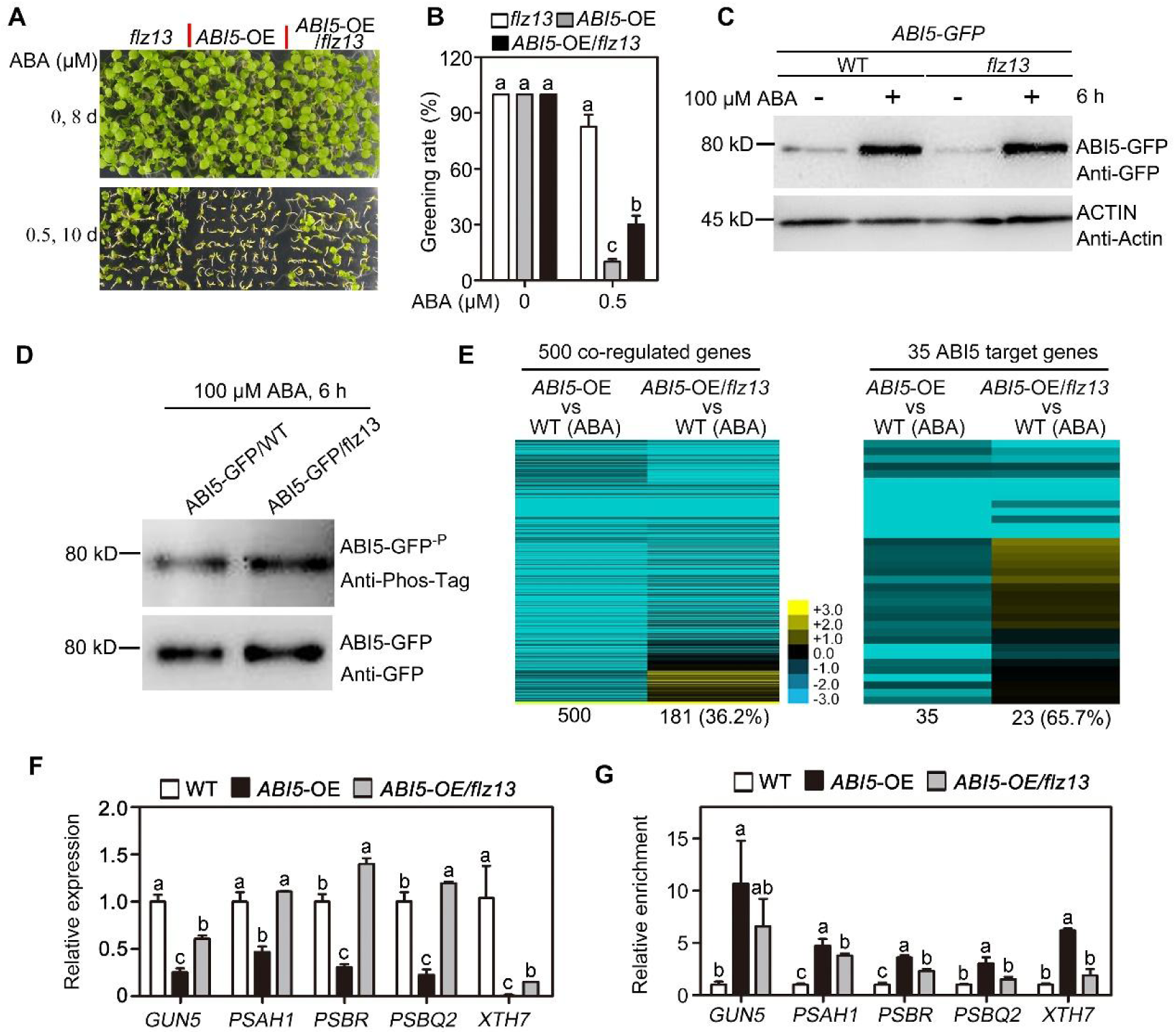
The function of ABI5 during seed germination is partially compromised in *FLZ13* knockdown mutants. A, Seedlings of *flz13*, *ABI5*-OE, and *ABI5*-OE/*flz13* germinated on the medium containing different concentrations of ABA for the indicated time points. B, Cotyledon greening of *flz13*, *ABI5*-OE, and *ABI5*-OE/*flz13* upon different concentration ABA treatment at the indicated time points. Data are presented as means ± SD (*n* = 3 biological replicates). For each biological replicate, around 80 seeds of each genotype were used for seedling establishment assay. The different letters above each bar indicate statistically significant differences as determined by a one-way ANOVA test followed by Tukey’s multiple test (P < 0.05). C, Immunoblotting analysis of ABI5-GFP protein in WT and *flz13* backgrounds. Whole seedlings of 7-day-old *ABI5-GFP* and *ABI5-GFP/flz13* were treated with or without 100 μM ABA for 6 h before protein extraction for western blotting analysis using Anti-GFP and Anti-Actin antibodies. D, Phosphorylation of ABI5-GFP protein in WT and *flz13* backgrounds upon ABA treatment. Whole seedlings of 7-day-old *ABI5-GFP* and *ABI5-GFP/flz13* were treated with 100 μM ABA for 6 h before protein extraction for immunoprecipitation using GFP-Trap beads, then the samples were subjected to Western blotting analysis with Anti-GFP and Anti-Phos-tag antibodies. E, Heat map showing the expression of ABA/ABI5/FLZ13 co-regulated genes in *ABI5*-OE and *ABI5*-OE/*flz13* lines in the presence of 0.5 μM ABA. 3-day-old germinating seeds with 0.5 μM ABA treatments were collected for RNA-seq assay. F, qRT-PCR assay showing the expression of the selected genes in WT, *ABI5*-OE and *ABI5*-OE/*flz13* upon 0.5 μM ABA treatment. 3-day-old germinating seeds with 0.5 μM ABA treatments were collected for qRT-PCR assay. *PP2A* was used as the internal control. Data are presented as means ± SD (*n* = 3 technical replicates). The different letters above each bar indicate statistically significant differences as determined by a one-way ANOVA test followed by Tukey’s multiple test (P < 0.05). This experiment repeated twice with similar results. G, ChIP-qPCR showing the relative enrichment of ABI5 to the selected gene locus. Data are presented as means ± SD (*n* = 3 technical replicates). The different letters above each bar indicate statistically significant differences as determined by a one-way ANOVA test followed by Tukey’s multiple test (P < 0.05). This experiment repeated twice with similar results.

We then analyzed the expression of 500 co-regulated genes in germinating seeds of *ABI5*-OE #1 *and ABI5*-OE #1/*flz13* upon ABA treatment by RNA-seq assay **(Supplemental Data 13)**. Heat map showed significantly altered expression level of 181 genes (36.2%) **(Figure 6E, leaf panel)**, indicating that ABI5-mediated transcription regulation of these genes is affected by FLZ13. For instance, 23 out of 35 ABI5 target genes showed differential expression between *ABI5*-OE #1 and *ABI5*-OE #1/*flz13* plants under ABA treatment **(Figure 6E, right panel)**. We also conducted qRT-PCR to confirm the transcript levels of *GUN5*, *PSAH1*, *PBSR*, *PBSQ2* and *XTH7* **(Figure 6F)**. Based on these results, we speculated that FLZ13 might affect the DNA binding ability of ABI5 to its downstream genes. To test this possibility, we performed ChIP-qPCR analysis using ABA-treated germinating seeds of *ABI5*-OE #1 and *ABI5*-OE #1/*flz13*. The obtained results showed that the enrichment of ABI5 on the promoters of *GUN5*, *PSAH1*, *PBSR*, *PBSQ2* and *XTH7* was generally decreased in *ABI5*-OE #1/*flz13* than that in *ABI5*-OE #1 **(Figure 6G)**, indicating that knockdown of *FLZ13* decreased the binding of ABI5 to its target genes in planta.

## Discussion

ABA modulates plant growth or stress response by triggering massive transcriptional reprogramming, which mainly depends on several ABA-responsive transcription factors, among them, ABI5 functions as the master regulator during ABA-inhibited seed germination and seedling establishment (Chen *et al*., 2020). However, how *ABI5* modulates ABA-repressed seed germination remains largely elusive, primarily because only few target genes of ABI5 are directly associated with germination process (Lee and Luan, 2012; Skubacz *et al*., 2016). Furthermore, the precise regulatory mechanism regarding to ABI5-engaged transcription regulation machinery also need to be further clarified.

In this study, we employed TurboID-mediated proximity labeling to obtain ABI5 interactome and identified FLZ13 as a new interacting protein of ABI5 **(Figure 1)**. FLZs are plant-specific regulatory proteins containing a common FLZ domain or DUF581 (Domain of Unknown Function 581) (Jamsheer and Laxmi, 2015). Two recent studies suggested that FLZ genes are involved in plant ABA response (Chen et al., 2021; Jamsheer et al., 2018). For instance, ectopic expression *ZmFLZ25* in Arabidopsis results in increased sensitivity of plants to ABA during seed germination (Chen *et al*., 2021), but the precise regulatory mechanism needs further characterization. Our current results not only demonstrated that FLZ13 positively regulates ABA signaling, but also uncovered the underlying molecular mechanism. Firstly, FLZ13 directly interacts with ABI5, a core transcription factor in ABA signaling (Skubacz *et al*., 2016), and this interaction is responsive to ABA treatment **(Figure 1; Supplemental Figure 4)**. Secondly, progenies of the *flz13* mutant were relatively less sensitive to ABA, while those of the *FLZ13* overexpressing plants were much more sensitive to ABA when compared to WT during seed germination and seedling establishment **(Figure 2)**. Thirdly, FLZ13 and ABI5 regulate common target genes in response to ABA **(Figure 4)**. Lastly, FLZ13 and ABI5 function interdependently to mediate ABA response during seed germination and seedling establishment **(Figure 5** and **6)**. These results strongly indicated that FLZ13 works along with ABI5 to positively regulate ABA signaling during seed germination. We also verified that ABA receptors and SnRK2 kinase are required for *FLZ13* to positively regulate ABA signaling **(Figure 2D** and **E)**.

Genetic analysis showed that ABI5 and FLZ13 work synergistically to positively regulate ABA signaling during seed germination **(Figure 3, 5 and 6)**. The *flz13-1 abi5* double mutant exhibited higher germination and greening percentages than *abi5* and *flz13-1* single mutants in the presence of ABA, while *FLZ13-OE* / *ABI5-OE* showed higher sensitivity to ABA than *FLZ13-OE* and *ABI5-OE* (**Figure 3**). Interestingly, the ABA sensitivity of the progenies of *flz13-1 ABI5-OE* were between *flz13-1* and *ABI5-OE* plants (**Figure 6A-C**), indicating that the action of ABI5 during ABA signaling partially requires FLZ13. FLZ13 does not affect the protein stability or phosphorylation of ABI5 in the presence of ABA **(Figure 6D** and **E)**, but partially impairs the DNA binding ability of ABI5 to downstream target genes **(Figure 6G)**. Given that the bZIP domain required for dimerization and DNA binding of ABI5 (Finkelstein and Lynch, 2000; Dröge-Laser et al., 2018) is involved in the interaction with FLZ13 **(Figure 1D)**, it is perhaps surprising that addition of FLZ13 enhances rather than inhibits ABI5 function. As FLZ13 is a zinc-finger protein and can be recruited to *GUN5*, *PBSR*, *PSAH1*, *PBSQ2* and *XTH7* promoters *in vivo* through interacting with ABI5 **(Figure 5H)**, it’s possible that the FLZ13-ABI5 complex may function similarly as the dimers of ABI5 and has increased binding activity to promoters of target genes, similar in the case of ABI5-PRR5 module (Yang et al., 2021). Detailed biochemical mechanisms underlying how FLZ13 protein synergizes with ABI5 to modulate downstream genes deserve further investigation.

RNA-seq analysis identified 567 ABA, ABI5 and FLZ13 co-regulated genes **(Figure 4A; Supplemental Data 8)**. Interestingly, most co-regulated genes of FLZ13 and ABI5 are repressed by ABA **(Figure 4B)**, suggesting that ABI5 and FLZ13 play important roles in ABA-mediated transcriptional repression. Interestingly, these co-repressed DEGs were mainly enriched in categories related to growth, including “photosynthesis” (41 genes, *p* value = 3.6E-22) and “cell wall organization” (40 genes, *p* value = 1.7E-11) **(Figure 4D; Supplemental Data 9)**. But as we know, ABI5 is a transcription factor with strong transcription activation activity, so how ABI5-FLZ13 complex represses the target gene expression remains an open question. It is possible that ABI5-FLZ13 complex might recruit transcriptional corepressors to target gene loci and modulate their expression. For example, the mediator (a bridge between transcription factors (TFs) and RNA polymerase II (RNA Pol II) during transcription) subunit MED16 are shown to interact with ABI5 to repress the expression of ABI5 target genes (Guo *et al*., 2021). In our ABI5-interactome data, several transcriptional corepressors such as LEUNIG, SUESS and TETRATRICOPEPTIDE REPEAT 2 (TPR2) (Franks et al., 2002; Long et al., 2006) showed significantly higher enrichment in ABI5-Turbo-GFP vs Turbo-GFP pair **(Supplemental Data 1)**, and they were positioned at the central nodes in the PPI networks **(Supplemental Figure 2)**, indicating the important roles of them in ABI5-mediated transcriptional repression. Further dissection of the underlying molecular mechanisms will provide new insights into the genetics basis of ABI5-FLZ13-inhibited seed germination and postgerminative growth.

In summary, our study shed new light on how ABI5-mediated ABA signaling that represses seed germination and seedling establishment **(Figure 7)**. Under suitable conditions, following seed germination, ABA level decreases and the downregulation of ABI5 activity releases its suppression, which results in normal seedling establishment. If adverse conditions are imposed to germinating seeds, elevated ABA activates ABI5 and the ABI5-FLZ13 module arrests germination or seedling establish process by suppressing growth-related genes till suitable conditions release this suppression and plant development restarts. These findings might be helpful for resetting the balance between growth and stress resistance based on ABA pathway, thus to engineer stress-resistant and high-yielding crops in future.

**Figure 7.**
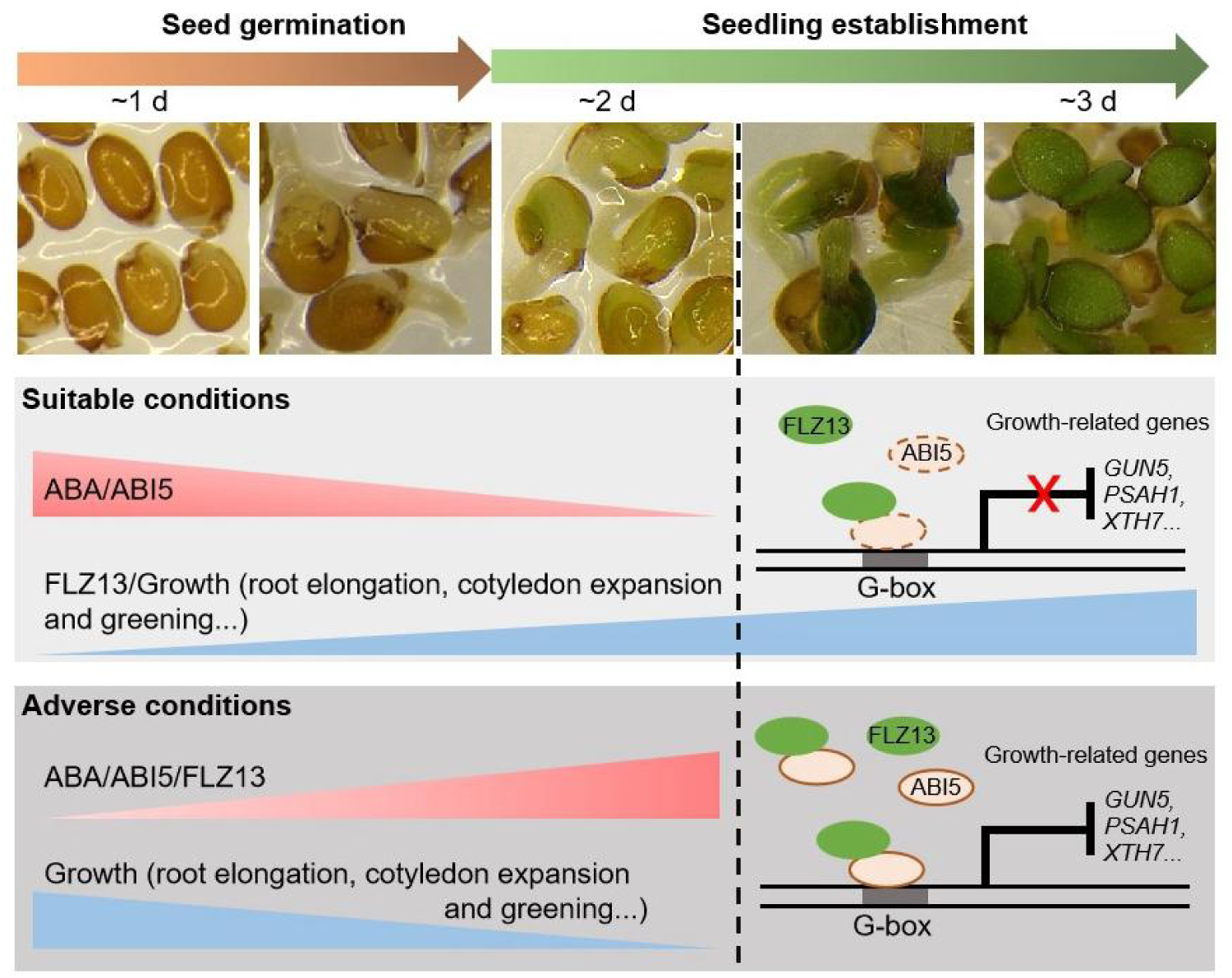
A working model for ABI5-FLZ13 module in seed germination and seedling establishment. Under suitable conditions, following seed germination, ABA level decreases and the downregulation of ABI5 activity releases its suppression, which results in normal seedling establishment. If adverse conditions are imposed to germinating seeds, elevated ABA activates ABI5 and the ABI5-FLZ13 module arrests germination or seedling establish process by suppressing growth-related genes till suitable conditions release this suppression and plant development restarts.

### Material and methods

#### Plant materials and growth conditions

All the Arabidopsis (*Arabidopsis thaliana*) lines used in this study were in the Col-0 background. The *abi5-8* mutant and *ABI5* overexpression plants (*pUBQ10::ABI5-GFP*, *ABI5*-OE #1) have been described previously (Li *et al*., 2019; Zhou *et al*., 2015). The *flz13* (SALK_142112C) mutant was obtained from ABRC (https://abrc.osu.edu/). To generate *FLZ13*-OE transgenic lines, full-length *FLZ13* was cloned into the binary vector pCAMBIA1300 in-frame fused with a *green fluorescence protein* (*GFP*) at the C-terminus under the control of *Arabidopsis UBQ10* promoter. The resulting construct was introduced into Col-0, *pry1pyl12458* (Gonzalez-Guzman *et al*., 2012) and *snrk2.2*/*2.3* (Fujii *et al*., 2007) by *Agrobacterial*-mediated transformation using flower dip method (Clough and Bent, 1998). To generate *pFLZ13::FLZ13-GFP* plants, the entire genomic sequence of *FLZ13* containing a ∼ 2.0 kb promoter sequence was PCR-amplified and inserted into the binary vector pCAMBIA1300 to in-frame fused with a *GFP* at the C-terminus. The resulting construct was introduced into Col-0 by *Agrobacterial*-mediated transformation using flower dip method. To generate *gFLZ13/flz13* (*gFLZ13-C*) lines, the entire genomic sequence of *FLZ13* containing a ∼ 2.0 kb promoter sequence and ∼1.0 kb terminator sequence was PCR-amplified and inserted into the binary vector pCAMBIA1300. The resulting construct was introduced into *flz13* by *Agrobacterial*-mediated transformation using flower dip method. Full-length *TurboID* was cloned into pCAMBIA1300 in-frame fused with a *GFP* at the C-terminus under the control of *Arabidopsis UBQ10* promoter to generate TurboID-GFP construct. To generate *ABI5-TurboID-GFP*, full-length *ABI5* was inserted into TurboID-GFP to fuse with TurboID-GFP at the C-terminus. The resulting constructs were introduced into Col-0 by *Agrobacterial*-mediated transformation using flower dip method. T_0_ seeds were screened in half-strength Murashige & Skoog (MS) medium containing hygromycin (25 mg/ml). Homozygous plants of the T3 generation were used for subsequent studies. The *abi5 flz13*, *FLZ13*-OE/*abi5* and *ABI5*-OE/*flz13* were generated by crossing and genotyping, respectively. The primers used were listed in **Supplemental Data 14**.

The seeds were surface sterilized with 30% (v/v) bleach for 5 min, washed with ddH_2_O for 3 times (3 min every time), then kept at 4°C in the darkness for 3 d before being sown on 1/2 MS plates supplemented with 0.8% (w/v) agar and 1% sucrose. After 6 d, the well-established seedlings were transferred to soil and grown in a chamber at 22°C under long-day conditions (LD, 16 h light/ 8 h dark). *Nicotiana Benthamian* plants were also grown in the same chambers under long-day conditions for BiFC and LCI assays.

### ABA sensitivity test

The germination and greening of seeds from different genotype were determined as described previously (Li *et al*., 2019). Briefly, seeds were surface sterilized, then cold stratified at 4°C/dark for 3 d before being sown on 1/2 MS medium plates with or without supplementation of ABA. The plates were placed in an artificial growth chamber at 22°C under LD conditions for germination. After 4 d, germination was determined based on the appearance of the embryonic axis (appearance of radicle protrusion) as observed under a microscope. After 6 to 12 d, pictures were taken to calculate the greening rate (appearance of green cotyledons on seedlings) with the photoshop software.

### Proximity labeling, affinity purification and mass spectrometry

Five-day-old seedlings of *pUBQ10::TurboID-GFP* (Tb-GFP #4) and *pUBQ10::ABI5-TurboID-GFP* (ABI5-Tb-GFP #3) transgenic seedlings grown on 1/2 MS media plates were transferred to 1/2 liquid medium supplemented with 50 μM of biotin for incubation for 1 h. Then, seedlings were rinsed with ice cold water tree times for 5 min each to stop the labeling reaction. The seedlings were then dried with paper towels, and split into aliquots of about 0.6 g fresh weight for the three biological replicates, and stored at −80°C until use.

Total protein was extracted from the collected samples with 6 mL buffer (50 mM Tris pH 7.5, 150 mM NaCl, 0.1% SDS, 1 mM EDTA, 1% Triton-X-100, 0.5% Na-deoxycholate, 1 mM DTT, 1x complete protease inhibitor cocktail, 1 mM PMSF). Before affinity purification with streptavidin-coated magnetic beads (MedChemExpress, Cat No.HY-K0208), the protein extract was subject to a desalting procedure to remove free biotin using Zeba™ Spin Desalting Columns (Thermofisher, Cat No.89893) according to the manufacturer’s instructions. AP was conducted at 4°C for about 16 h. After affinity purification, samples on beads were washed five times with the extraction buffer, then 1% samples were eluted for immunodetection with Streptavidin-HRP (GeneScript, Cat No.M000-91) and the remaining samples were digested on-beads with trypsin, and analyzed by LC–MS/MS following our established protocols as previously described (Branon et al., 2018; Mair *et al*., 2019).

### Proteomic analysis and protein-protein interaction (PPI) network construction

A detailed proteomic analysis procedure has been described previously (Mair *et al*., 2019). Filtering and statistical analysis were done with Perseus (Tyanova et al., 2016). The output file from MaxQuant was imported into Perseus using the LFQ intensities as Main category. The data matrix was filtered to remove proteins marked as ‘only identified by site’, ‘reverse’ and ‘potential contaminant’. LFQ values were log2 transformed and proteins that were identified/quantified only in one of three replicates of one genotype were removed. The criteria for identification of ABI5-specific substrates were fold change (ABI5-Turbo-GFP vs TurboID-GFP) > 2, *P* < 0.05. Finally, the volplot was generated on the OmicShare tools platform (http://www.omicshare.com/tools) and the PPI were constructed on line (https://metascape.org/; (Zhou et al., 2019).

### Protein interaction analysis

(1) Yeast-two hybrid (Y2H) assay. The full-length and truncated versions of *FLZ13* or others tested proteins were cloned into pGBKT7 to generate bait vectors with Gal4 DNA binding domain (BD). The primers used were listed in **Supplemental Data 14**. The full-length and truncated versions of *ABI5* were cloned into pGADT7 to generate prey vectors with Gal4 activation domain (AD) as described previously (Li *et al*., 2019). Yeast two-hybrid assays were performed as described previously (Yang et al., 2018). The bait and prey vectors were cotransformed into the yeast strain AH109 and physical interactions were determined by the growth of co-transfected yeast cells on synthetic dropout medium (SD)-Leu-Trp or SD-Leu-Trp-His-Ade plates for 3-4 d after plating.

(2) Bimolecular fluorescence complementation assay (BiFC). The full-length coding sequence of *FLZ13* and *ABI5* were cloned into the pCAMBIA1300-YN or pCAMBIA1300-YC vectors to generate FLZ13-YN and ABI5-YC, respectively. The primers used were listed in **Supplemental Data 14**. Then, the resulting constructs were transformed into *Agrobacterium tumefaciens* GV3101. *A. tumefaciens* containing FLZ13-YN, *A. tumefaciens* containing ABI5-YC, and *A. tumefaciens* containing Histon3-mCherry were co-injected into fully-expanded leaves of 5-week-old *N. Benthamian* plants at a ratio of 2: 2: 1. After infiltration for 3 d, the infiltrated regions were observed and imaged with a confocal microscope.

(3) Luciferase complementation imaging assays (LCI). The full-length coding sequence of *FLZ13* and *ABI5* were cloned into the pCAMBIA1300-nLUC and pCAMBIA1300-cLUC vectors to generate FLZ13-nLUC and cLUC-ABI5, respectively. The primers used were listed in **Supplemental Data 14**. Then, the resulting constructs were transformed into *Agrobacterium tumefaciens* GV3101. *A. tumefaciens* containing FLZ13-nLUC, *A. tumefaciens* containing cLUC-ABI5, and *A. tumefaciens* containing P19 were co-injected into fully-expanded leaves of 5-week-old *N. Benthamian* plants at a ratio of 2: 2: 1. After infiltration for 3 d, the infiltrated regions were injected with 1 mM of fluciferin before capturing LUC activities with Tanon-5200. For ABA treatment, the leaves were pre-treated with 10 μM of ABA for 4 h.

(4) Pull-down assay. The full-length *FLZ1*3 and *ABI*5 were cloned into pMAL-c2× and pGEX4T-3, respectively. The primers used were listed in **Supplemental Data 14**. Then, the recombinant vectors were transformed into *E. coli* BL21 (DE3) to express His-MBP-FLZ13 and GST-ABI5 proteins at 25°C for 6 h induction with 0.4 mM IPTG. The supernatant containing His-MBP or His-MBP-FLZ13 proteins was mixed with 50 μl MBP-magic beads (Sangon) and incubated for 2 h at 4 °C. Then, the beads were washed six times with 1 ml PBS and incubated with the supernatant containing GST-ABI5 protein for 4 h at 4 °C. After washing six times with 1 ml PBS, the beads were boiled in 100 μl 1x SDS-PAGE loading buffer. Finally, the presence of GST-tagged and MBP-tagged proteins was detected by Western blotting using anti-His and anti-GST antibodies, respectively.

### Gene expression analysis by quantitative real-time PCR (qRT-PCR)

To analyze the gene expression of *FLZ13* and *ABI5* during seed germination and their response to ABA, 50 mg Col-0 seeds were surface sterilized, then cold stratified at 4°C/dark for 3 d before being sown on 1/2 MS medium plates with or without supplementation of 0.5 μM ABA. The plates were placed in an artificial growth chamber at 22°C under LD conditions for germination and the seeds were harvested at the indicated time points for total RNA isolation using a Hipure plant RNA Mini kit (Magen). Total RNA (1 μg) was reverse-transcribed using HiScript^Ⓡ^ ⅡQ Select RT SuperMix (Vazyme), and the resulting first strand cDNA was used as a template for quantitative real-time PCR, which was performed in CFX96 Touch Real-Time PCR Detection System (Bio-Rad) and iTaq^TM^ Universal SYBR^Ⓡ^ Green Supermix (Bio-Rad). The expression of *PP2A* or *Actin2* was used as an internal control and the relative expression was calculated using the 2^-△△ CT^ method. The primers for qRT-PCR are listed in **Supplemental Data 14**.

### RNA Sequencing and Data Analysis

To perform RNA-sequencing analysis, 200 mg seeds were surface sterilized, then cold stratified at 4°C/dark for 3 d before being sown on 1/2 MS medium plates with or without supplementation of 0.5 μM ABA. The plates were placed in an artificial growth chamber at 22°C under LD conditions for germination. After 3 d, the seeds were harvested to isolate total RNA using a Hipure plant RNA Mini kit (Magen). RNA-seq was performed in Biomarker Technologies. Genes with Log2 (absolute fold changes) ≥ 1 and the averaged RPKM value from three biological repeats were higher than 1 at least from one pair were identified as reliable deferentially expressed genes (DEGs). Multiple testing was corrected via false discovery rate estimation and FDR below 0.01 were considered to indicate differential expression. The transcriptome data have been deposited in the NCBI GEO repository (http://www.ncbi.nlm.nih.gov/geo) with the accession no. PRJNA885244. The GO enrichment analysis was performed by using DAVID program (https://david.ncifcrf.gov/home.jsp) (Huang et al., 2009). The heat maps were generated in Cluster 3 software and visualized in Treeview.

### ChIP-quantitative PCR (ChIP-qPCR) assay

ChIP assays were performed as previously described (Gendrel et al., 2005; Yang et al., 2020). Briefly, 3 d-old germinating seeds with 0.5 μM ABA treatment of wild type, *FLZ13*-OE #1, *FLZ13*-OE #1/*abi5*, *ABI5*-OE #1 and *ABI5*-OE#1/*flz13* were cross-linked with 1% formaldehyde. Then, the chromatin was extracted and sheared to ∼ 500 base pair by sonication before being immunoprecipitated with Magic-GFP trap (Chromotek). After cross-linking was reversed, the amount of each precipitated DNA fragment was determined by quantitative PCR. The relative quantity value was calculated by the 2^-△△CT^ method and presented as the relative ratio of IP to Input. The primers used in ChIP-qPCR assay were listed in **Supplemental Data 14**.

### Electrophoretic mobility shift assay (EMSA)

The EMSA experiment was performed as previously described (Li et al., 2022b). Briefly, full-length ABI5 and FLZ13 were separately fused with a His-MBP tag at the C terminal to generate the His-MBP-ABI5 and His-MBP-FLZ13 vectors, respectively. Then the resulting vectors were transformed into *E. coli BL21* (DE3) strains to express the recombination proteins at 25°C for 6 h induction with 0.4 mM IPTG. The expressed proteins were purified using His-magic beads (Sangon) from the soluble fractions. The biotin-labeled DNA probes are purchased from Sangon Biotech and listed in **Supplemental Data 14**. EMSA assay was performed using non-radioactive electrophoretic mobility shift assay Kit (Viagene Biotech, Ning Bo) following the instruction manuals.

### Protein detection and phosphorylation analysis

To analyze the protein levels of FLZ13 and ABI5 during seed germination and their response to ABA, 100 seeds of *pFLZ13::FLZ13-GFP* #9 were surface sterilized, then cold stratified at 4°C/dark for 3 d before being sown on 1/2 MS medium plates with or without supplementation of 0.5 μM ABA. The plates were placed in an artificial growth chamber at 22°C under LD conditions for germination and the seeds were harvested at the indicated time points. To check the protein level of ABI5, 7-d-old *ABI5*-OE #1 and *ABI5*-OE #1*/flz13* were subjected to 100 μM ABA for 6 h before being harvested. Total protein was extracted from the harvested samples using 20 mM Tris-HCl, pH 7.5, 150 mM NaCl, 1 mM EDTA, 1% trionx-100, and protease inhibitor cocktail tables (Roche Diagnostics).

For phosphorylation analysis of ABI5, 7-d-old *ABI5*-OE #1 and *ABI5*-OE #1*/flz13* were subjected to 100 μM ABA for 6 h before being harvested to isolate protein. The ABI5-GFP proteins were purified by GFP-trap then subjected to phosphorylation analysis using a Phos-tag Biotin BTL-111 kit.

### Data processing and statistical analysis

The histograms were constructed by GraphPad Prism software (http://www.graphpad.com) with the average values and standard deviation, and the original data were given in **Supplemental Data 15.** The unprocessed images for Western blots were give in **Supplemental Figure 9**. Statistical differences were calculated by one-way ANOVA in SPSS program (https://spssau.com/) or Student’s *t*-test in Microsoft Excel. Different letters indicate means that were statistically different by Tukey’s multiple testing methods (P < 0.05). * P < 0.05, ** P < 0.01.

## Author contribution

C.Y., C.G., Y. W., and M.L. conceived the project. C.Y., X.L., L.Y., X.C., K.L., J.L., C.L., X.Z., and H. L. performed the experiments. C.Y., C.G., Y.L., S.Z., X.Z., P.R., Y.W., and M.L. analyzed the results. C.Y., C.G., Y.L., X.Z., P.R., Y. W., and M.L. wrote or edited the manuscript. All authors read and approved of its content.

---

**Supplemental Figure 1:**
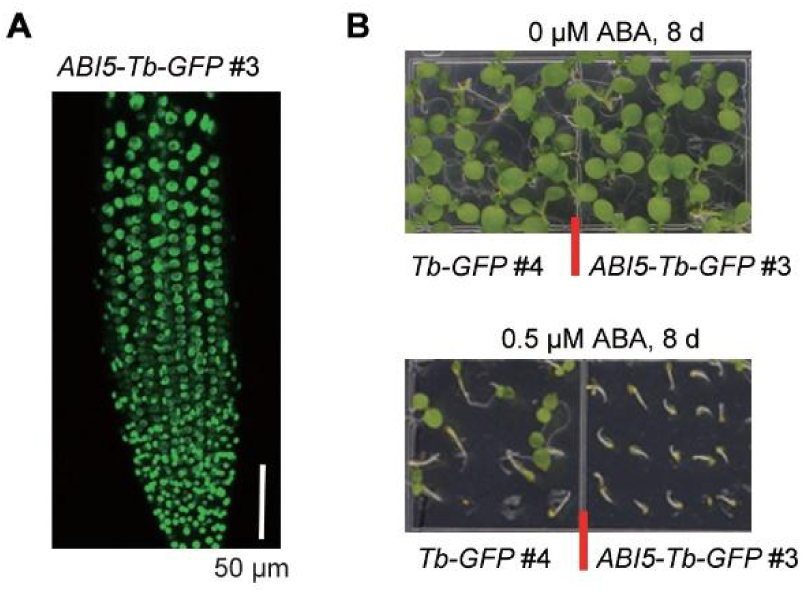
Characterization of Tb-GFP and ABI5-Tb-GFP transgenic plants. A, Confocal image showing the nuclear-localization of ABI5-Tb-GFP proteins in root cells of 5-day-old seedlings. B, Seedlings of Tb-GFP #4 and ABI5-Tb-GFP #3 upon 0 or 0.5 μM ABA treatment for 8 d.

**Supplemental Figure 2:**
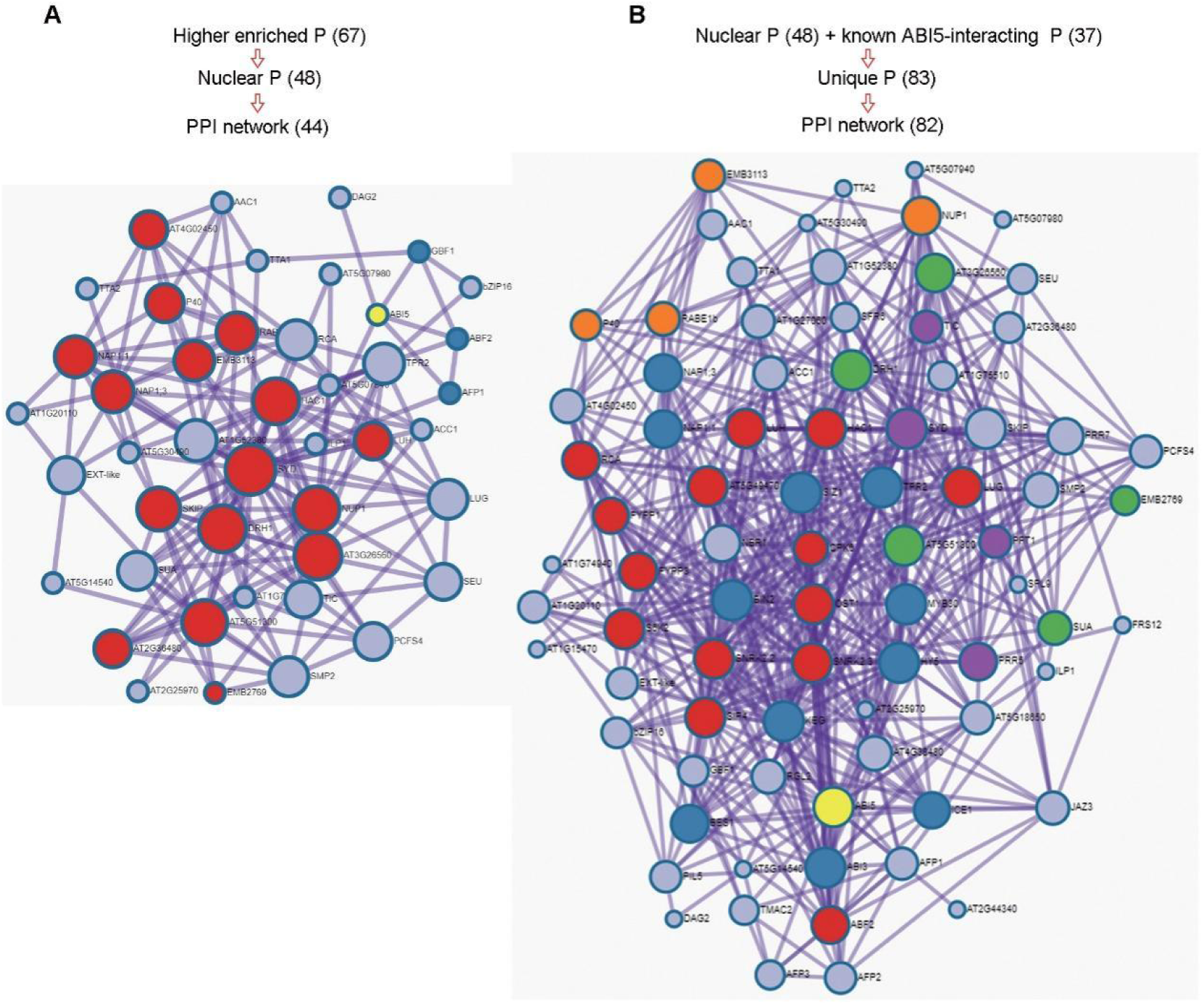
PPI of ABI5 and its interacting proteins. ABI5 is highlighted in yellow. The PPI were generated on line (https://metascape.org/; Zhou et al., 2019). P, protein.

**Supplemental Figure 3:**
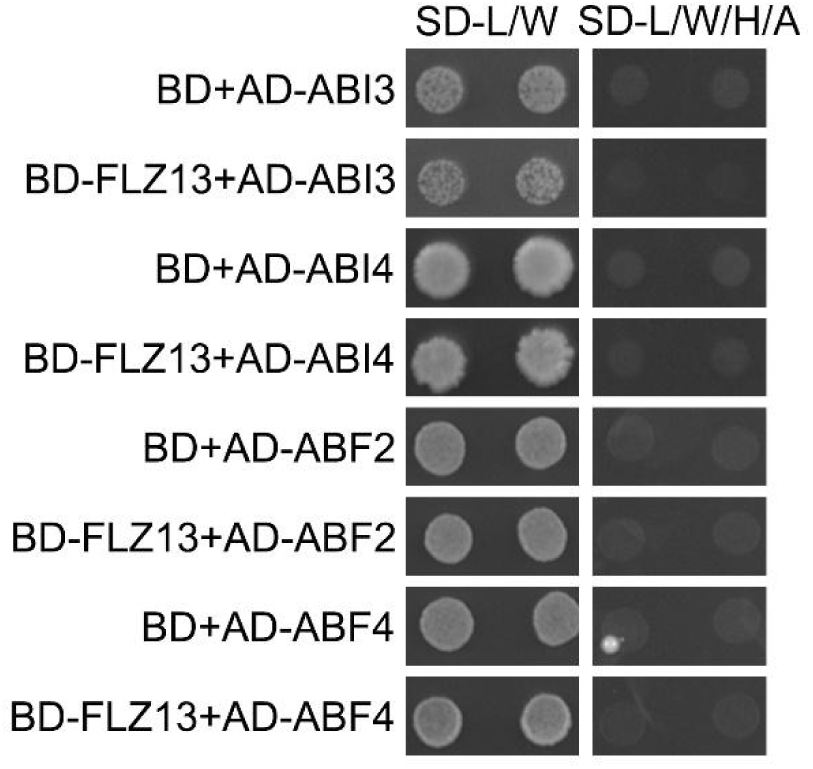
Y2H analysis of the interaction between FLZ13 and other key transcription factors involved in ABA signaling. Protein interaction was determined by growth of the yeast cells co-transformed with various combinations of the plasmids on synthetic dropout medium lacking Leu and Trp (SD-LW) and synthetic dropout medium lacking Leu, Trp, His, and adenine (SD-LWHA).

**Supplemental Figure 4:**
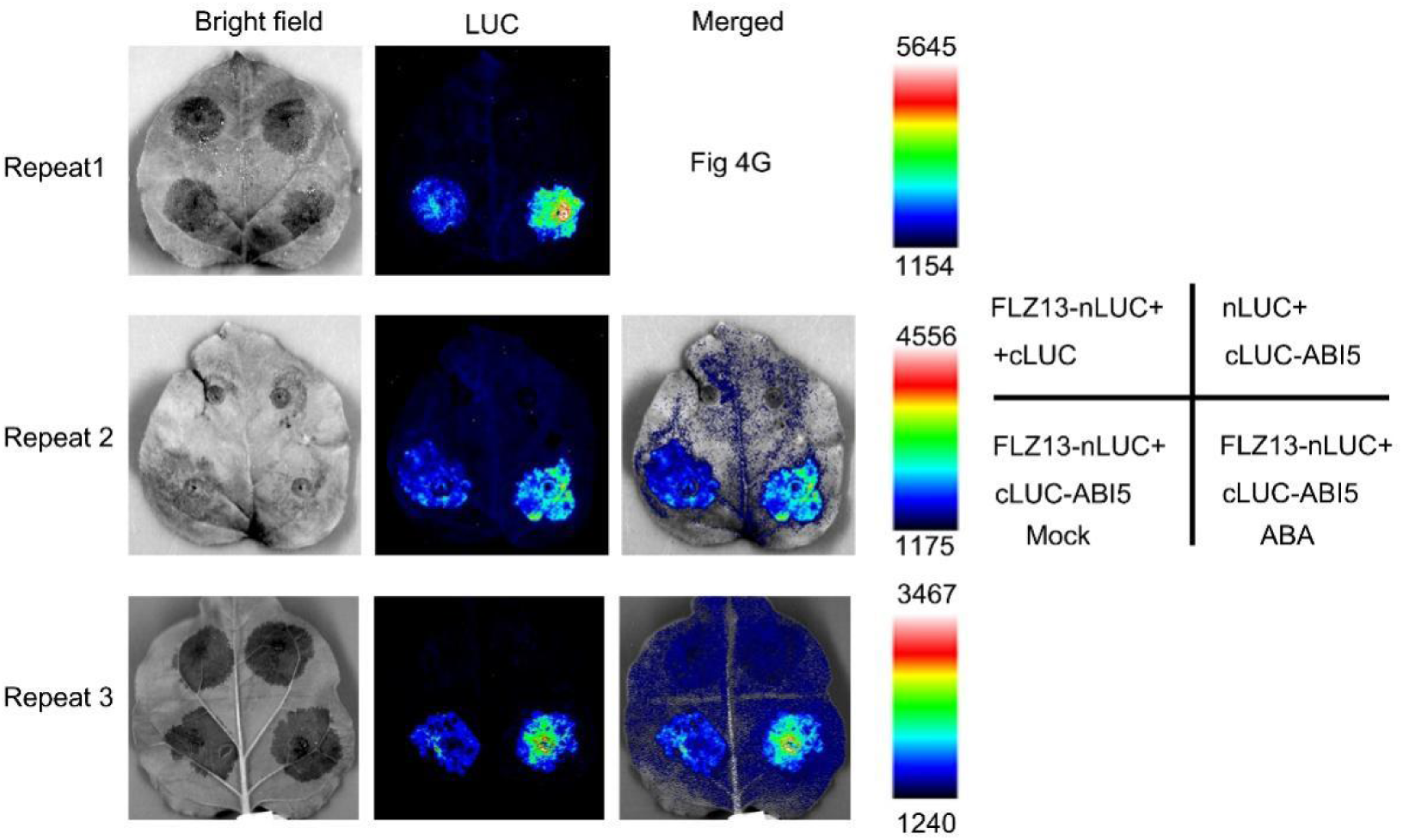
More biological replications of LCI experiments showing ABA promotes FLZ13-ABI5 interaction. Three independent experiments of LCI showing the interaction between FLZ13 and ABI5, and the promoting effects of ABA on this interaction.

**Supplemental Figure 5:**
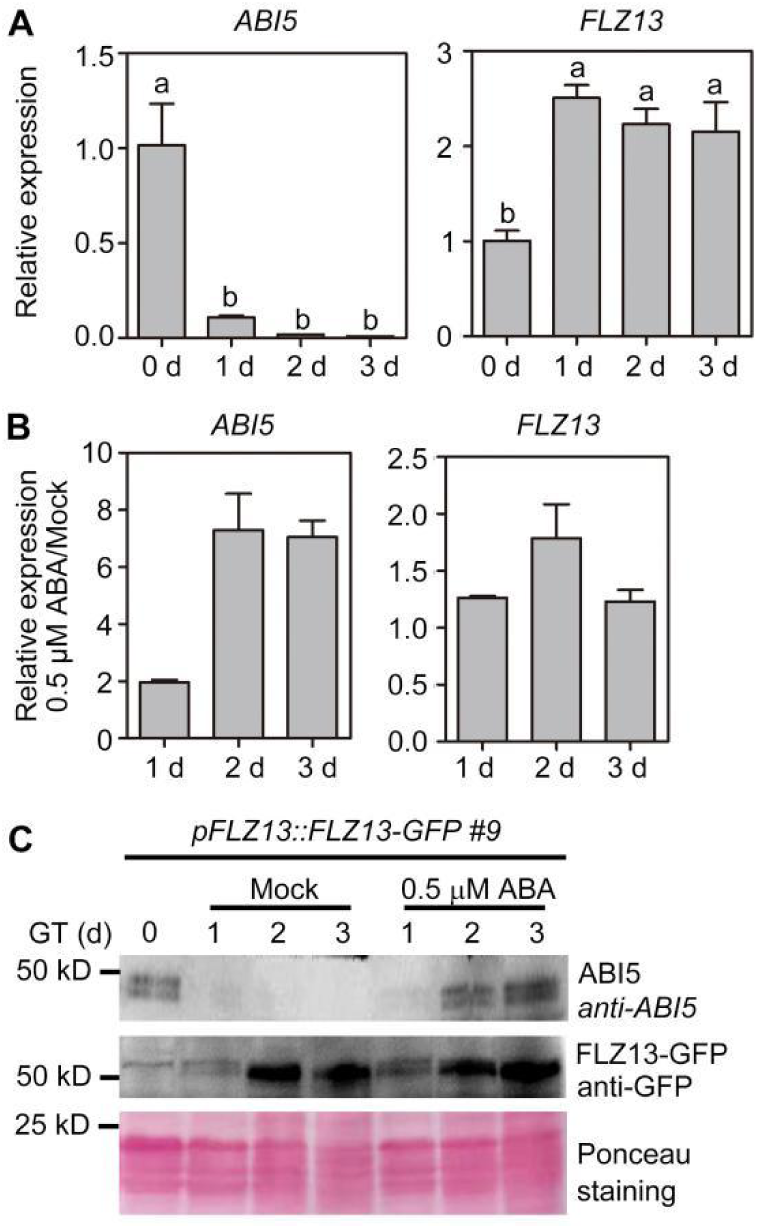
Expression pattern of *FLZ13* and *ABI5* during seed germination with/without ABA treatment. A, qRT-PCR analysis of FLZ13 and ABI5 expression in dry and germinating seeds. Actin2 was used as the internal control. B, qRT-PCR analysis of FLZ13 and ABI5 expression in response to ABA during seed germination. The data were presented as the ratio of ABA to Mock treatment. Actin2 was used as the internal control. C, Immunoblot analysis of FLZ13 and ABI5 protein levels in response to ABA during seed germination. GT: germination time. Data presented in A and B are mean ± SD, n = 3. Statistically significant differences between means were calculated by one-way analysis of variance. Different letters in A indicate statistically significant differences between means by Tukey’s multiple comparison test (P < 0.05).

**Supplemental Figure 6:**
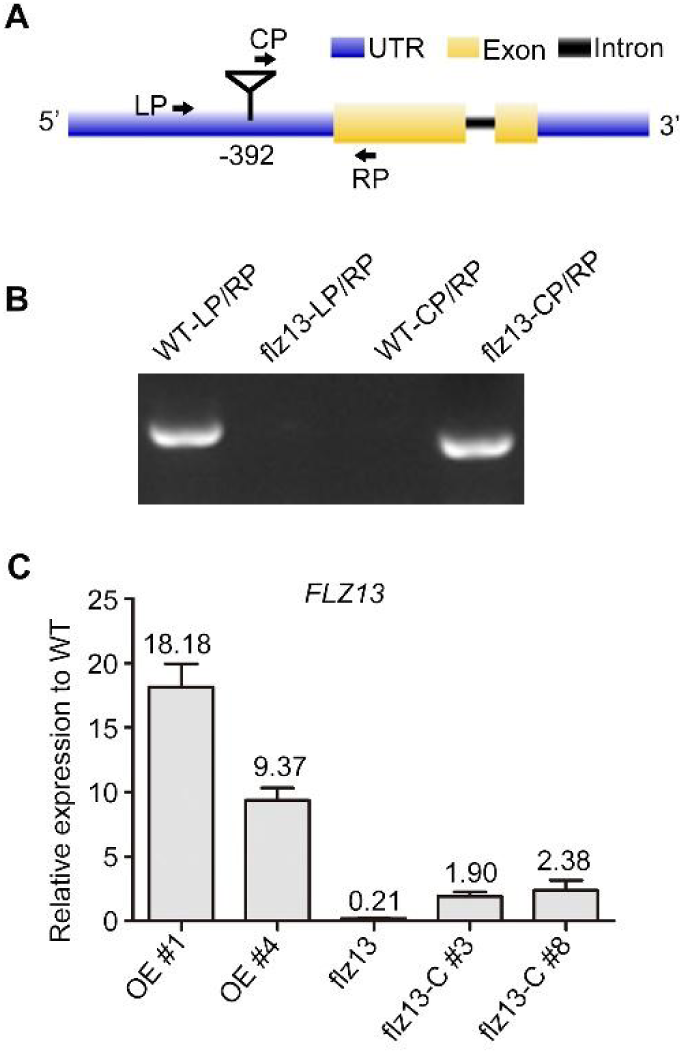
Characterization of *FLZ13*-related plant materials. A, Schematic gene structure of FLZ13. Triangle indicates the T-DNA insertion site in flz13 mutant. Arrows indicate primers used for genotyping of flz13 mutant. B, Gel electrophoresis of genotyping PCR with WT and flz13 DNA using primers in A. C, qRT-PCR shows the gene expression of FLZ13 in the indicated genetic background. Data is mean ± SD, n = 3. Actin2 was used as the internal control. The expression level in WT was set as 1.0. The numbers on the bars are the fold change.

**Supplemental Figure 7:**
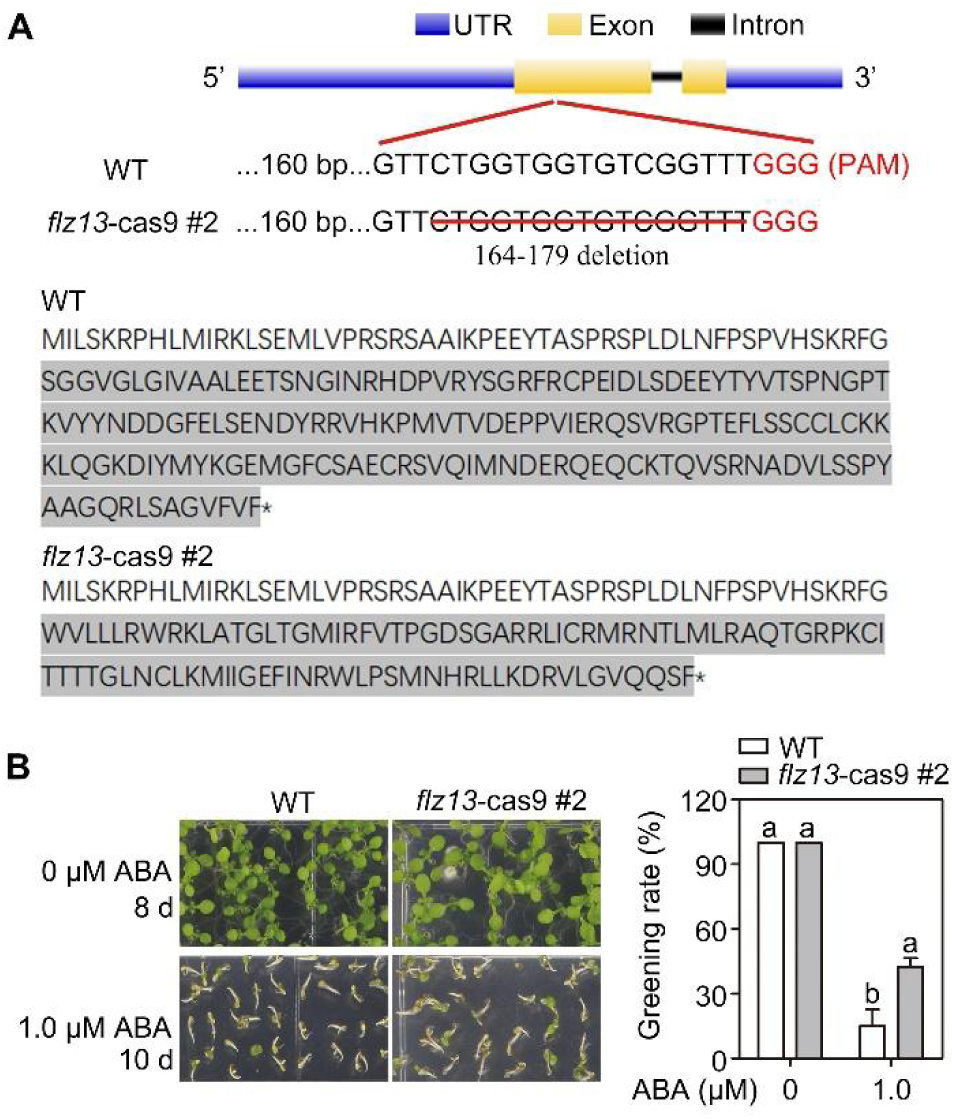
Characterization of *flz13*-cas9 plants. A, The CRISPR/Cas9 mutants of *FLZ13*. *flz13*-cas9 #2 harbors a 15-nucleotides deletion which is predicted to encode a truncated protein. B, Seedlings and greening rate of WT and *flz13*-cas9 #2 upon 0 or 1.0 μM ABA treatment as recorded at the indicated time points. Data is mean ± SD, *n* = 3. Statistically significant differences between means were calculated by one-way analysis of variance. Different letters indicate statistically significant differences between means by Tukey’s multiple comparison test (*P* < 0.05).

**Supplemental Figure 8:**
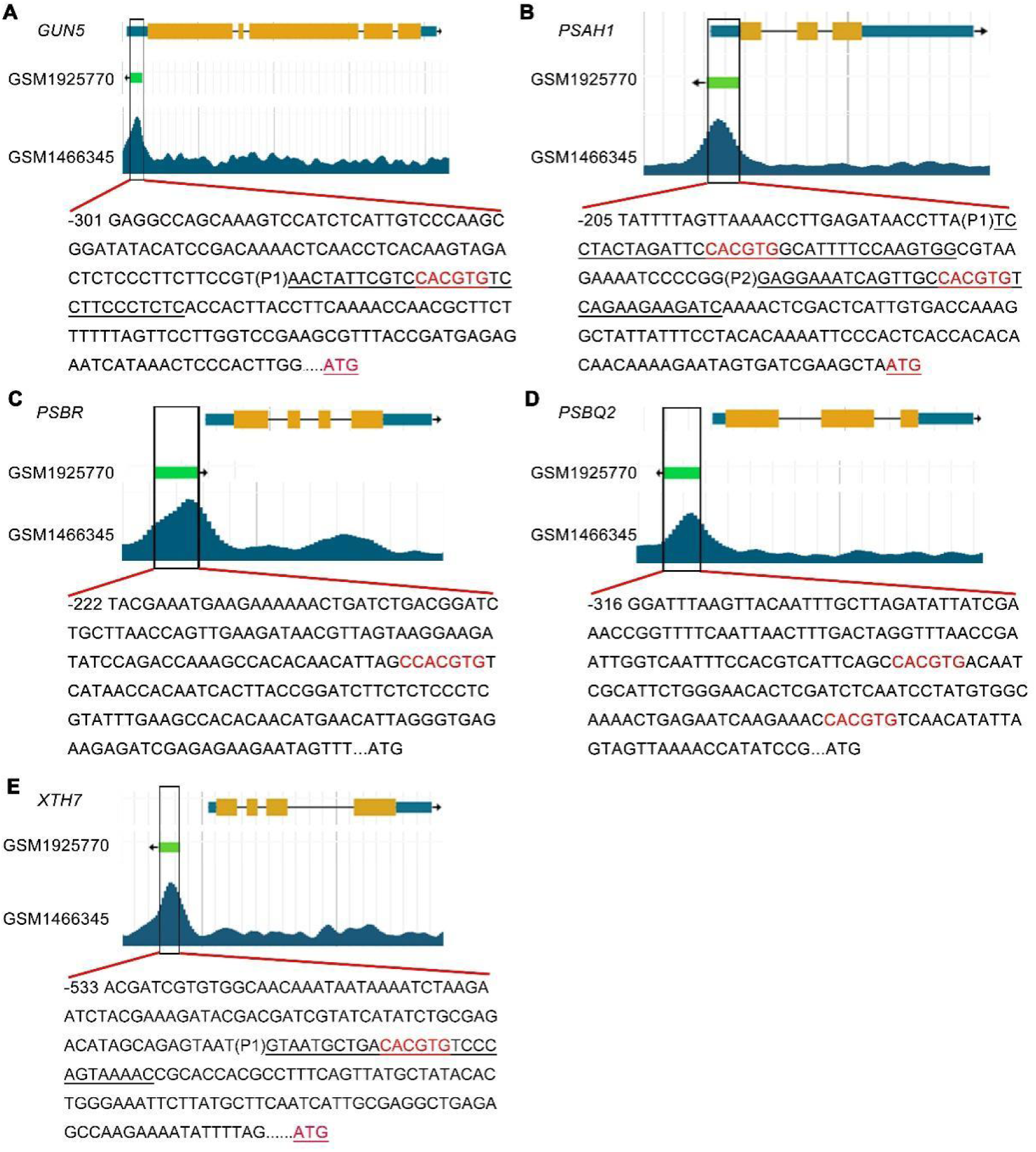
Five selected target genes of ABI5. The raw data were downloaded from http://bioinfo.sibs.ac.cn/plant-regulomics/ (Ran et al., 2020) and edited in photoshop. The G-box and primers for EMSA analysis were underlined.

**Supplemental Figure 9:**
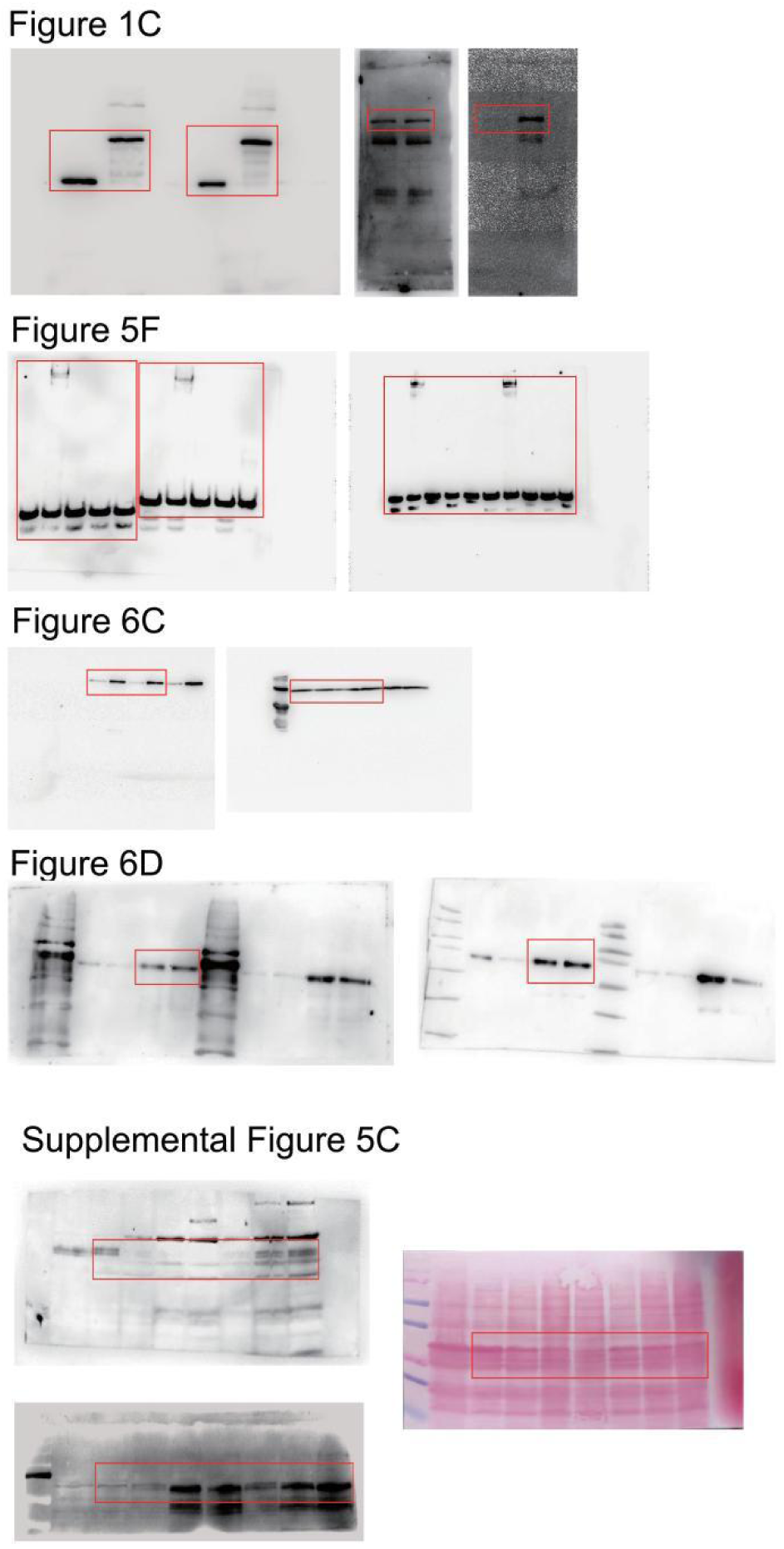
Original images for Western blots. The red boxes marked the sheared regions presented in the corresponding figures.

## Supplemental Data

The following materials are available in the uploaded Excel file.

**Supplemental Data 1**: List of the higher enriched proteins in ABI5-Tb-GFP vs Tb-GFP comparison.

**Supplemental Data 2**: List of the 48 nuclear-localized proteins of 67 identified higher enriched proteins in Supplemental Data 1.

**Supplemental Data 3**: List of the 37 known ABI5-interacting proteins.

**Supplemental Data 4**: DEGs in response to ABA treatment in WT germinating seeds.

**Supplemental Data 5**: Expression of known ABA-induced marker genes in the RNA-seq data.

**Supplemental Data 6**: DEGs in ABI5-OE seeds compared to WT upon ABA treatment.

**Supplemental Data 7**: DEGs in FLZ13-OE seeds compared to WT upon ABA treatment.

**Supplemental Data 8**: Expression of 567 ABA, FLZ13 and ABI5 co-regulated genes.

**Supplemental Data 9**: GO enrichment of 496 ABA/ABI5/FLZ13 co-repressed genes.

**Supplemental Data 10**: Target genes of ABI5.

**Supplemental Data 11**: List of the 35 ABI5 target genes co-regulated by ABA-ABI5-FLZ13.

**Supplemental Data 12**: Expression of the 500 ABI5/FLZ13 co-regulated genes in WT, *FLZ13-OE* and *FLZ13-OE*/*abi5* plants upon ABA treatment.

**Supplemental Data 13**: Expression of the 500 ABI5/FLZ13 co-regulated genes in WT, *ABI5-OE* and *ABI5-OE/flz13* plants upon ABA treatment.

**Supplemental Data 14**: List of primers used in this study.

**Supplemental Data 15**: Original data for statistical analysis.

